# The Role of BAZ2-dependent Chromatin Remodeling in Suppressing G4 DNA Structures and Associated Genomic Instability

**DOI:** 10.1101/2024.12.10.627505

**Authors:** Adrianna L. Vandeuren, Kierney O’Dare, Rosemary H. C. Wilson, Patrick van Eijk, Lindsay R. Julio, Shannon G. MacLeod, Ella Chee, Annika Salpukas, Emma M. Kriz, George A. Lantz, Shellaina Gordon, Simon J. Elsässer, Simon H. Reed, Tovah A. Day

## Abstract

DNA G-quadruplexes (G4s) are secondary structures with significant roles in regulating genome function and stability. Dysregulation of the dynamic formation of G4s is linked to genomic instability and disease, but the underlying mechanisms are not fully understood. In this study, we conducted a screen of chromatin-modifying enzymes and identified nine potential inhibitors of G4 formation, including seven that were not previously characterized. Among these, we highlight the role of BAZ2 chromatin remodelers as key suppressors of G4 DNA and G4-related genome instability. Depletion of BAZ2 subunits led to increased G4 formation, especially at transcriptional regulatory elements. BAZ2B was found to associate with G4 loci, suggesting that it plays a direct role in suppressing G4s. While BAZ2-deficient cells exhibited modest genomic instability, treatment with the G4-stabilizing ligand BRACO19 exacerbated double-strand breaks (DSBs), highlighting its utility as a tool to study G4-dependent genome instability. DSB profiling using INDUCE-seq uncovered distinct breakage patterns around G4s, further underscoring the impact of G4s on genome integrity. Notably, we found that within G4s, G repeats were more susceptible to DSBs than loops. These results establish BAZ2 chromatin remodeling complexes as direct regulators of G4 dynamics and provide new insights into G4-dependent genome instability.

## Introduction

DNA can adopt secondary structures that affect genome function and contribute to genome instability. Among these, the G-quadruplex (G4) features four DNA strands that create stacked tetrads of Hoogsteen-bonded guanines, stabilized by a central cation. With known roles in transcription (1–4), replication (5, 6), DNA repair (7), telomere maintenance (8), distal chromosome interactions (9, 10), and contributions to genomic instability (11), G4 structures play key roles in cellular function. Mapping their genomic locations is therefore fundamental to fully understanding these roles. Computational algorithms predict over 3 million G4 motifs in the human genome (12), with approximately 700,000 sequences forming G4 structures in naked DNA (13, 14). Yet, studies with G4-structure specific antibodies, such as BG4 (15), detect far fewer G4 structures in the chromatin context (16–19) and reveal that G4 structures vary by cell type (1, 19, 20), hinting at a regulatory role for chromatin environment. Indeed, histone acetylation is positively correlated with G4 structure formation, while DNA methylation has a negative correlation (19, 21, 22). Together, these observations emphasize the influence of chromatin environment in G4 structure regulation, yet the full extent of this G4 regulation has not been defined.

Improper regulation of G4 structures increases the risk of genomic instability, with evidence from multiple genomes indicating that G4s can trigger DNA double-strand breaks (DSBs) and mutations. In the yeast and *C. elegans* genomes, G4-forming sequences are linked to spontaneous DSBs (23–25) which are more common in the absence of specific DNA repair factors (24–29). In human cells, G4 motifs are associated with chromosomal rearrangements, as evidenced by their enrichment near breakpoints of recurrent tumor rearrangements (22, 30–33). Consistent with this observation, DSBCapture analysis of endogenous lesions in a transformed cell line also showed enrichment for G4-forming sequences (34). Cell permeable G4 ligands that stabilize G4 structures may offer a useful approach for studying DSBs linked to G4 formation. Structurally diverse ligands like pyridostatin (PDS), CX-5461, and BRACO19 induce DSBs in human cells (35–37), with PDS and CX-5461 specifically reported to target G4-enriched loci (35, 36); however, the role of BRACO19 at these loci has not yet been assessed. Although G4 structures are strongly linked to genomic instability, the exact mechanisms by which they induce DSBs are not yet fully elucidated.

BAZ2A and BAZ2B are paralogs of the Bromodomain Adjacent to Zinc finger domain 2 (BAZ2) family which is part of the Imitation Switch (ISWI) remodeler family. They function as regulatory subunits that partner with either the SMARCA1 or SMARCA5 catalytic subunits (38, 39). Interestingly, SMARCA5 interacts biochemically with the G4 structure in the c-Myc promoter (40). With 39% identity and 63% similarity, the BAZ2 paralogs are capable of repositioning, inserting, or evicting nucleosomes on DNA (38). BAZ2 remodeling complexes appear to influence diverse cellular phenotypes by remodeling regulatory elements to drive transcriptional changes. BAZ2A, for example, is overexpressed in several cancers and is associated with poor prognosis (41, 42). As part of the nucleolar remodeling complex (NoRC), it fosters heterochromatin formation at ribosomal DNA loci (43), leading to decreased transcription of both rDNA and additional genes (44). Conversely, BAZ2 inhibition in mice enhances liver regeneration by upregulating protein synthesis pathways (45), and baz-2 deletion in *C. elegans* has been linked to lifespan extension and greater stress resilience mediated by transcriptional changes (46). The potential involvement of G4 structures in BAZ2-dependent chromatin remodeling has not been previously explored.

In this study, we identify a collection of chromatin alterations that regulate G4 DNA structure formation. Among them, we report the first identification of a chromatin remodeling complex, BAZ2, that suppresses the formation of G4 DNA. Inhibition or depletion of either BAZ2A or BAZ2B led to global increases in G4 DNA, with G4s emerging across various genomic elements. We present evidence that BAZ2A and BAZ2B have both distinct and overlapping roles in preventing G4 formation. We demonstrate that BAZ2B occupancy overlaps substantially with G4 structures, suggesting a direct mechanism of suppression. Depletion of either BAZ2 subunit, in combination with G4 ligand stabilization, induces DSBs, hallmarks of genomic instability. We use INDUCE-seq to accurately map DNA DSBs genome-wide and observe an enrichment of G4 structures in proximity to DSBs. In addition, we find that the G4 ligand BRACO19 induces genomic instability, which may be exacerbated by BAZ2A depletion. On a kilobase scale, BRACO19 generates specific DSB patterns surrounding G4 structures, while closer analysis reveals that within G4 motifs, G repeats are more susceptible to DSBs than loop regions. Our data suggest that BRACO19 promotes genome instability at G4 sites, making it an effective tool for exploring G4-dependent DSB patterns. Overall, our findings align with a model in which BAZ2-mediated chromatin remodeling restricts G4 DNA structure formation, and loss of this activity triggers G4-dependent genomic instability.

## Results

### A screen identifies chromatin modifications that control G4 DNA formation

To identify chromatin modifications that promote or suppress G4 DNA formation, we used immunofluorescence (IF) microscopy to screen a library of 253 small molecule inhibitors of chromatin modifying enzymes (**Supplementary Fig. 1A and Supplementary Table 1**). We measured the nuclear accumulation of G4 DNA using BG4, a structure-specific antibody that detects G4s (15), and identified 28 inhibitors that increased global G4 DNA to a similar or greater extent than pyridostatin (PDS), a G4 ligand (35) (**Fig. 1A**). To address the possibility that any increases in G4 DNA during prolonged inhibitor treatment were due to cell cycle perturbations (15), we re-screened these 28 hits with a shorter 2-hour treatment, minimizing potential cell cycle impacts. Using this shorter duration of inhibitor treatment, we confirmed that 14 out of the 28 inhibitors increased global G4 DNA (**Supplementary Fig. 1B**). We noticed that several putative hits inhibit histone deacetylation including PTACH, Tacedinaline, and RGFP966 (47–49) (**Table 1**, **Fig. 1B**). Previous studies have reported that increases in histone acetylation lead to global increases in G4 structures (21, 50), indicating that our screen had identified chromatin modifications in line with established findings. We therefore focused on the remaining 11 inhibitors that target processes other than histone acetylation (**Table 1**).

**Figure 1:**
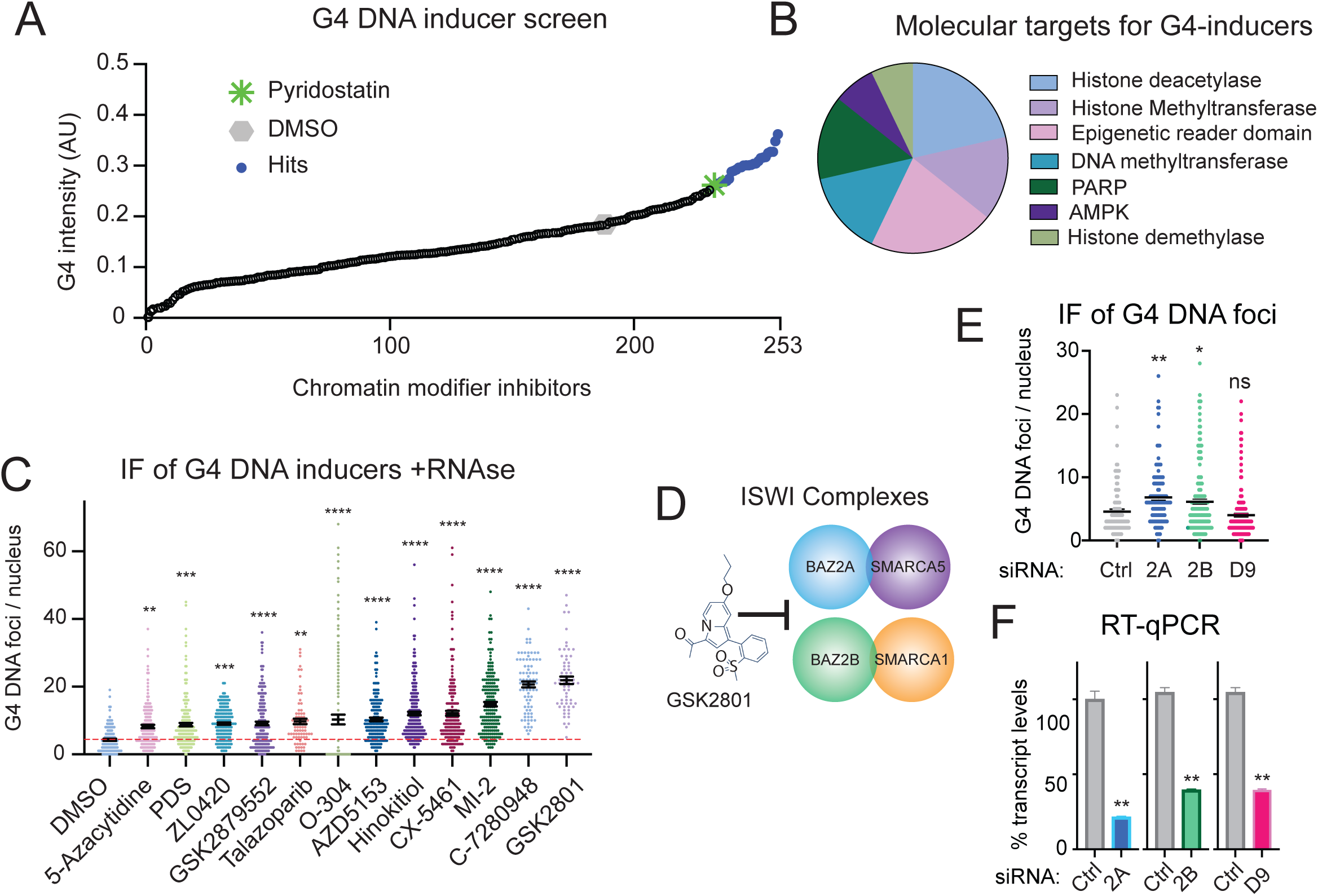
Identification of chromatin modifiers that suppress G4 DNA formation. **A.** Quantitation of mean immunofluorescence intensity of BG4 following 24 hr treatment with 10 µM of a library of small molecule inhibitors of chromatin modifiers or G4 ligands. The negative control, DMSO, is labeled in grey, pyridostatin (PDS) in green, and hits with an intensity similar or greater than PDS in blue. **B**. Targets of the inhibitors identified as hits. AMPK, adenosine monophosphate-activated protein kinase. PARP, poly (ADP-ribose) polymerase. **C.** Quantitation of BG4 DNA foci following a 2 hr inhibitor treatment (10 µM) with RNAse A. The red dashed line corresponds to the mean for DMSO-treated cells. **D.** Schematic representation of GSK2801 targeting BAZ2-containing ISWI complexes. **E.** Quantitation of G4 DNA foci per nucleus with the indicated siRNA at 48 hr. **F.** Quantitation of transcript levels measured by RT-qPCR normalized to β-actin. Ctrl, Control; 2A, BAZ2A; 2B, BAZ2B; D9, BRD9. ns, non-significant; *, *p* < 0.05; **, *p* < 0.01; ***, *p* < 0.001; ****, *p* < 0.0001 (One-way ANOVA for multiple comparisons).

**Table 1.**

| **Inhibitor** | **Enzyme target** | **Epigenetic modification** |
| --- | --- | --- |
| **Hinokitiol, 5-Azacytidine** | DNMT1 | DNA methylation |
| **O-304** | AMPK | Phosphorylation |
| **GSK2879552** | LSD1 | Histone methylation |
| **AZD5153, ZL0420** | BRD4 | Binds acetyl lysine |
| **Talazoparib tosylate, Veliparib** | PARP | Poly(ADP)ribosylation |
| **C-7280948** | PRMT1 | Histone demethylation |
| **MI-2** | MEN1/MLL | Multiple roles |
| **PTACH, Tacedinaline, RGFP966** | HDACs | Histone deacetylation |
| **GSK2801** | BAZ2 | Heterochromatin |

We verified by IF with RNase treatment that these 11 inhibitors led to increases specifically in G4 DNA, and not G4 RNA (**Figure 1C, Supplementary Fig. 1C**), as BG4 is known to bind G4 RNA as well (51). The 11 validated inhibitors targeted eight distinct enzymatic activities, with DNMT1, BRD4, and PARP1 each represented by two inhibitors among our identified hits (**Table 1**). Of these hits, we focused our attention on GSK2801, an inhibitor of the Bromodomain Adjacent to Zinc finger domain 2 (BAZ2) protein family (52) (**Fig. 1D, Supplementary Fig. 2A**) that has not been previously reported to influence G4 DNA formation.

### BAZ2 chromatin remodeling complexes suppress G4 DNA formation

GSK2801 inhibits BAZ2A and BAZ2B, while also showing cross-reactivity with BRD9 (52). Using siRNA, we found that depletions of either BAZ2A or BAZ2B, but not BRD9, led to significant increases in G4 DNA formation (**Fig. 1E-F, Supplementary Fig. 2B-C**). We also verified that neither treatment with GSK2801 nor depletion of BAZ2A or BAZ2B caused significant cell cycle perturbations (**Supplementary Fig. 2D-E**). Taken together, these data indicate that abrogation of BAZ2, whether by chemical inhibition or specific depletion, leads to a significant increase in global G4 DNA in human cells.

To identify the specific G4 structures that arise in the absence of BAZ2, we performed Cleavage Under Targets and Tagmentation (CUT&Tag) for G4 DNA using the BG4 antibody (20, 53) with optional depletion of BAZ2A and BAZ2B by siRNA (**Fig. 2A**). High quality peaks were defined as those occurring in at least two of four replicates (**Fig. 2B, Supplementary Fig. 3A**). Consistent with our IF results, we identified a larger number of G4 peaks in either the siBAZ2A or siBAZ2B conditions relative to siControl using MACS2 (54) (**Fig. 2B-C**). To address the possibility of variation in CUT&Tag efficiencies across samples, we re-analyzed the data using SEACR (**Supplementary Fig. 3B**), a strategy optimized for CUT&RUN (55), and scaled the data to the number of *Drosophila* spike-in reads in each sample (**Supplementary Fig. 3C**). In this scaled analysis, siBAZ2A and siBAZ2B also led to a larger number of G4 peaks (**Supplementary Fig. 3B**).

**Figure 2:**
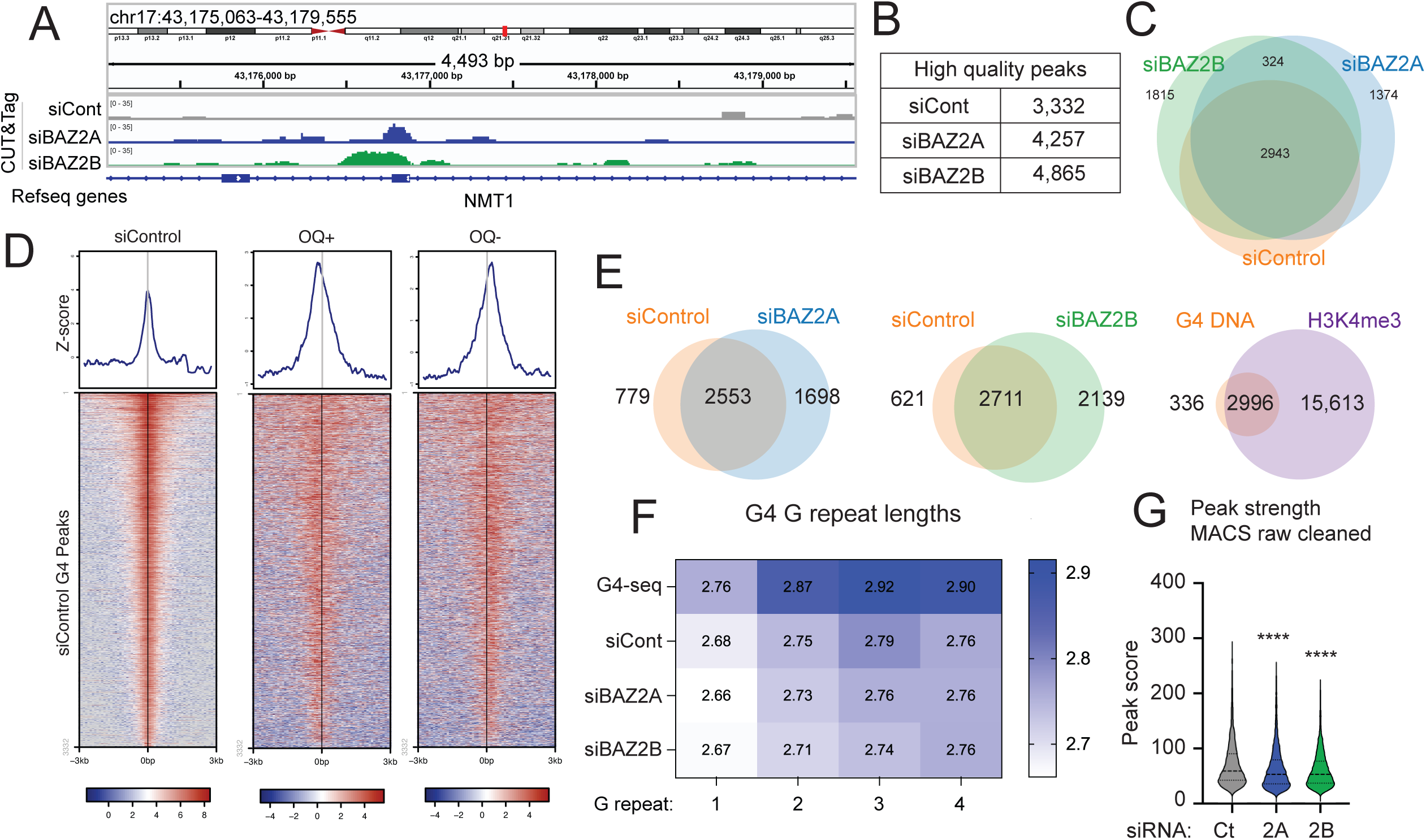
Characterization of G4 DNA in the absence of BAZ2. **A.** Genome browser view of G4 CUT&Tag signal at an example locus. **B.** Number of high quality G4 peaks in the indicated conditions 48 hr following siRNA. **C.** Overlap of G4 peaks between siRNA conditions. **D**. G4 CUT&Tag and observed quadruplexes (OQ) with PDS (1) as log2 transformed density heatmaps for peaks classified as G4 CUT&Tag in siControl (*n* = 3,332)**. E.** Venn diagrams showing overlap between G4 and H3K4me3 peaks with indicated siRNA. **F.** Mean of G repeat lengths of the first four G-runs identified (5’ to 3’ orientation) in G4 motifs within G4 CUT&Tag and G4-seq peaks (1). **G.** Peak score distribution for G4 CUT&Tag peaks with indicated siRNA. siCont, siControl; Ct, Control; 2A, BAZ2A; 2B, BAZ2B. (****, *p* < 0.0001; one-way ANOVA for multiple comparisons).

To examine the overall efficacy of the G4 CUT&Tag, we evaluated the number of G4 motifs identified by G4Catchall (56) in each MACS2 peak normalized to its length. We found a higher G4 motif density in G4 CUT&Tag peaks than in either a random size-matched control or in H3K4me3 peaks captured in parallel (**Supplementary Fig. 4A**), confirming that BG4 enriches for G4-containing loci. MEME motif discovery (57) analysis of MACS2 G4 peaks revealed high confidence, G-rich motifs consistent with G4 formation (**Supplementary Fig. 4B**). The G4 peaks in all three CUT&Tag conditions exhibited good overlap with observed quadruplexes (OQs) identified by G4-seq (**Fig. 2D, Supplementary Fig. 5A-C**), a whole genome sequencing strategy that exploits polymerase stalling at structured G4s (14). Under siBAZ2A treatment, 1,698 new G4 peaks were identified that were absent in siControl, whereas siBAZ2B produced 2,139 additional G4 peaks (**Fig. 2C**). While 324 peaks (∼15-19%) were common to both depletions, the rest were unique, suggesting that the two regulatory subunits have both common and distinct roles in G4 control. Therefore, BG4 CUT&Tag captured peaks that contain G4-forming sequences and depletion of either BAZ2 subunit led to an increase of both unique and shared G4s.

### BAZ2A- and BAZ2B-dependent G4 structures resemble those that form under basal conditions

We considered that BAZ2-dependent G4s might represent a distinct subset of G4s defined by genomic locations or structural features. First, we defined BAZ2A- and BAZ2B-dependent peaks as those that uniquely arise upon depletion of either factor (**Fig. 2E**); this comprises sets of 1,698 and 2,139 G4 structures for BAZ2A and BAZ2B, respectively. We classified the G4 peak subsets based on proximity to defined genomic elements using the Roadmap Epigenomics data (58). While H3K4me3 peak distributions varied relative to G4 peaks (**Supplementary Fig. 6A**), all three subsets of G4 peaks exhibited similar distributions, indicating that their proximity to specific genome elements is not a distinguishing factor. Next, we examined structural features of the G4 motifs detected by G4Catchall within our CUT&Tag peaks. The G-repeat lengths in these motifs were significantly shorter compared to those in the G4-seq dataset (*p* < 0.0001, Two-way ANOVA, **Fig. 2F**), but no other significant, consistent differences were found. Similarly, we did not observe consistent differences in either the loop length or loop composition (**Supplementary Fig. 6B-C**) although we noted an overall trend that the first loop was longer than the subsequent 3’ loops (*p* < 0.0001, Two-way ANOVA, **Supplementary Fig. 6B**). G4Hunter scores (12) for the G4s in each dataset revealed a modest but significant decrease in scores in the BAZ2B-dependent subset relative to siControl (*p* = 0.0016, Kolmogorov-Smirnov test, **Supplementary Fig. 6D**). Consistent with this trend, the mean peak intensity of G4 CUT&Tag peaks was significantly lower in both siBAZ2A and siBAZ2B (**Fig. 2G**), suggesting lower stability and greater propensity to convert to the canonical B form of DNA.

A recent study presented a framework for clustering G4s into families based on similar sequence, structure, and thermodynamic properties (59) and we wondered whether this multifactorial approach might help us further define the subset of BAZ2-dependent G4s. We clustered all the G4s with three tetrads identified in our CUT&Tag peaks into 95 pre-defined G4 families. Families 7, 37, 52, 79, and 80 were the most highly represented, comprising ∼43% of all 3-tetrad G4s across all three conditions (**Supplementary Fig. 7A-B**), with no obvious sequence similarities between these most highly represented G4 families. Overall, these analyses suggest that BAZ2-dependent G4s exhibit a modest tendency toward lower stability compared to G4s formed in unperturbed cells. However, in all other examined parameters, BAZ2-suppressed G4s closely resemble basal G4s.

### Integrative analysis of BAZ2B-containing remodeling complexes

We focused on BAZ2B in subsequent experiments as its cellular roles are less defined. To examine whether BAZ2B-containing chromatin remodeling complexes suppress G4 formation by directly binding to these loci, we sought to profile the landscape of endogenous BAZ2B occupancy in human cells. As none of the commercially available antibodies that we tested could detect endogenous BAZ2B, we used CRISPR/Cas9-mediated homology directed repair (HDR) to insert an HA tag at both endogenous alleles of BAZ2B in A549 cells (**Supplementary Fig. 8A-B**). We first verified that cells with HA-BAZ2B exhibited higher anti-HA IF signal than the parental cells (**Supplementary Fig. 8C-D**). Next, we performed CUT&Tag for HA-BAZ2B in A549 cells with optional siRNA-mediated BAZ2B depletion (**Supplementary Fig. 8E**) and observed that peak intensity for peaks common between siControl and siBAZ2B was significantly reduced in siBAZ2B-treated cells (**Supplementary Fig. 8F**), suggesting that HA-BAZ2B peaks represent sites of BAZ2B occupancy. HA CUT&Tag in cells containing the endogenous HA-BAZ2B tag identified 5,043 high quality peaks (present in at least two out of three replicates (**Supplementary Fig. 8G**)) that were absent in the parental control (**Fig. 3A-C**). MEME Suite analysis of HA-BAZ2B peaks identified multiple G-rich sequences, consistent with the potential for G4 formation (**Fig. 3D**). We noted that HA-BAZ2B peaks strongly corresponded with siBAZ2B G4 CUT&Tag peaks (**Fig. 3E**), whereas HA CUT&Tag peaks in parental cells, which lack an HA tag, did not exhibit the same pattern as they are likely background (**Supplementary Fig. 9A**). Indeed, the overlap between HA-BAZ2B and BAZ2B-dependent G4 peaks was statistically significant (GAT analysis (*p* = 0.001, 1,000 permutations) **Fig. 3F**) and 135-fold greater than would be expected by chance (60). Even those HA-BAZ2B peaks that did not overlap with G4 CUT&Tag peaks were enriched for G4-forming motifs (7.3 G4 motifs / 1,000 bp compared to 1.3 G4 motifs in a size-matched random control (**Supplementary Fig. 9B**)). Overall, these data suggest that BAZ2B colocalizes with G4-rich loci.

**Figure 3:**
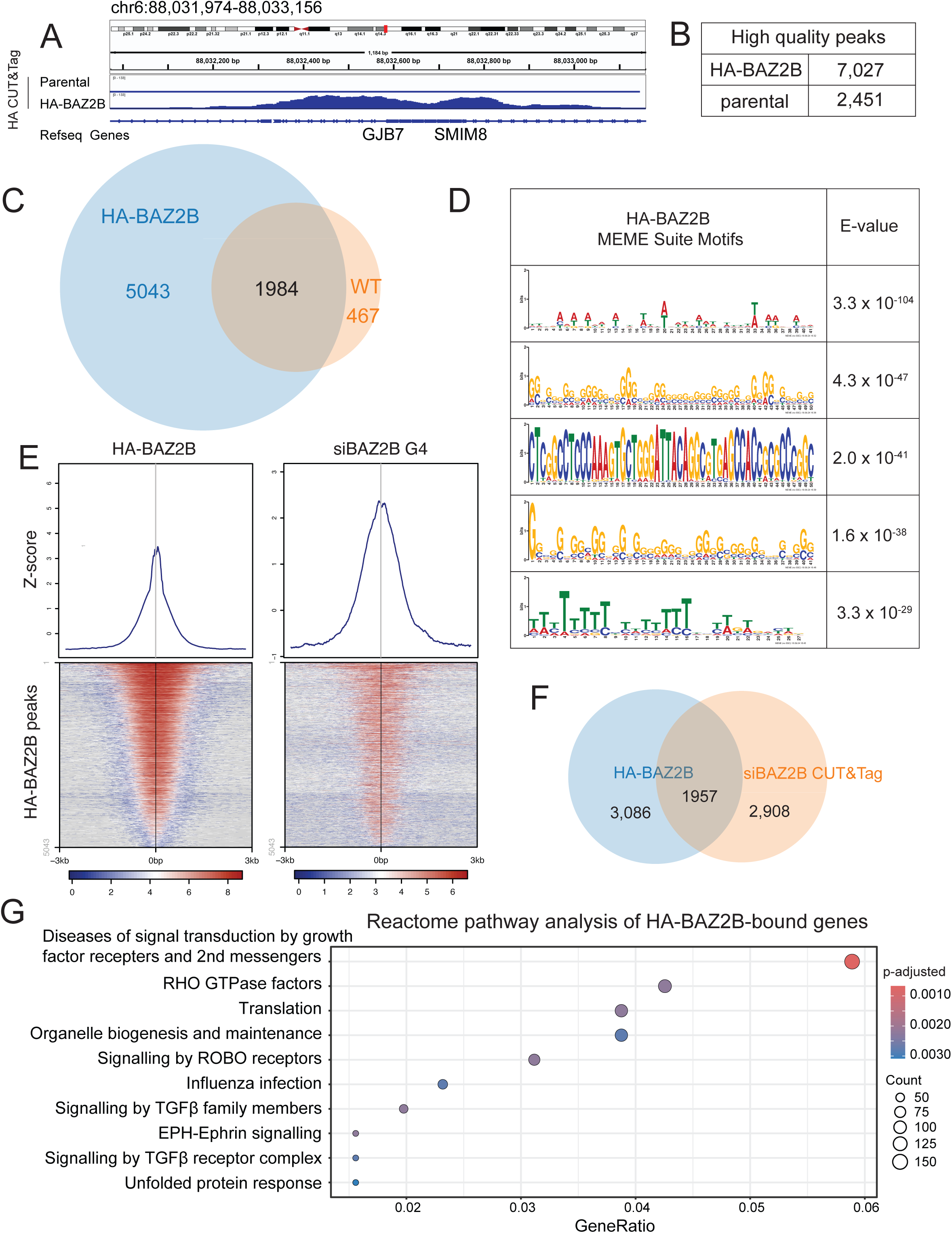
Genome-wide occupancy of endogenous BAZ2B. **A.** Genome browser view of HA CUT&Tag signal at an example locus. **B**. Number of high-quality HA peaks. **C**. Overlap between HA peaks in cells engineered with HA-BAZ2B endogenous tag and parental A549 cells that lack an HA tag. **D.** Motif discovery using MEME suite on HA-BAZ2B bound loci. **E.** HA-BAZ2B CUT&Tag and G4 CUT&Tag with siBAZ2B overlap in log2 transformed density heatmaps for peaks classified as HA-BAZ2B (*n* = 5,043)**. F.** Venn diagram of overlap of HA-BAZ2B peaks (*n* = 5,043) and G4 CUT&Tag peaks in siBAZ2B A549 cells (*n* = 4,865). **G.** Reactome pathway enrichment of HA-BAZ2B bound loci associated with TSSs (defined as within 3 kb of TSS).

Analysis of regulatory elements associated with HA-BAZ2B peaks showed that the majority correspond to active transcription start sites (TssAs, ∼31%) and promoters (∼54%). This pattern is similar to the G4s captured upon siBAZ2B and differs from random distributions (**Supplementary Fig. 9C**). In genic regions, HA-BAZ2B peaks show a striking accumulation at human TssAs (**Supplementary Fig. 9D**), and a similar phenomenon was observed in murine liver cells with ectopic expression of Flag-Baz2b ((45) **Supplementary Fig. 9E**). We performed Reactome pathway analysis (61) on genes within 3 kb of HA-BAZ2B peaks to capture both TssAs and promoters (**Fig. 3G**). Consistent with recently published functional data on murine Baz2b (45), we identified translation among the most significantly enriched pathways. Indeed, in human cells, we identified many BAZ2B-bound translation-associated genes including RPL22, RPL24, and RPLP1, whose homologs were Baz2b-regulated in the mouse study (45). Collectively, these findings suggest that BAZ2B suppresses the formation of G4 structures by directly binding to the G4-containing loci and has a conserved role of regulating translation in both human and murine cells.

### BAZ2 depletion leads to genome instability

We wondered whether elevated levels of G4 DNA in the absence of BAZ2 might result in increased genome instability. While BAZ2 depletion alone did not change cell cycle dynamics (**Supplementary Fig. 2E**), the combination of siBAZ2B with the G4 ligand BRACO19 (62) led to a significant increase in the sub-G1 fraction (**Fig. 4A, Supplementary Fig. 10A**), indicating potential apoptosis. We hypothesized that BRACO19 stabilizes the larger number of G4s in BAZ2-depleted cells, causing DSBs which could induce apoptosis. Consistent with this, we observed a significant increase in γH2AX foci, a marker of DSBs, in cells treated with both siBAZ2B and BRACO19 (**Fig. 4B, Supplementary Fig. 10B**), with more modest increases in γH2AX when BRACO19 was combined with siControl or siBAZ2A (**Fig. 4B**). Collectively, these findings suggest that the depletion of BAZ2 chromatin remodelers makes cells more vulnerable to BRACO19-induced genome instability, potentially indicating G4-dependent instability.

**Figure 4:**
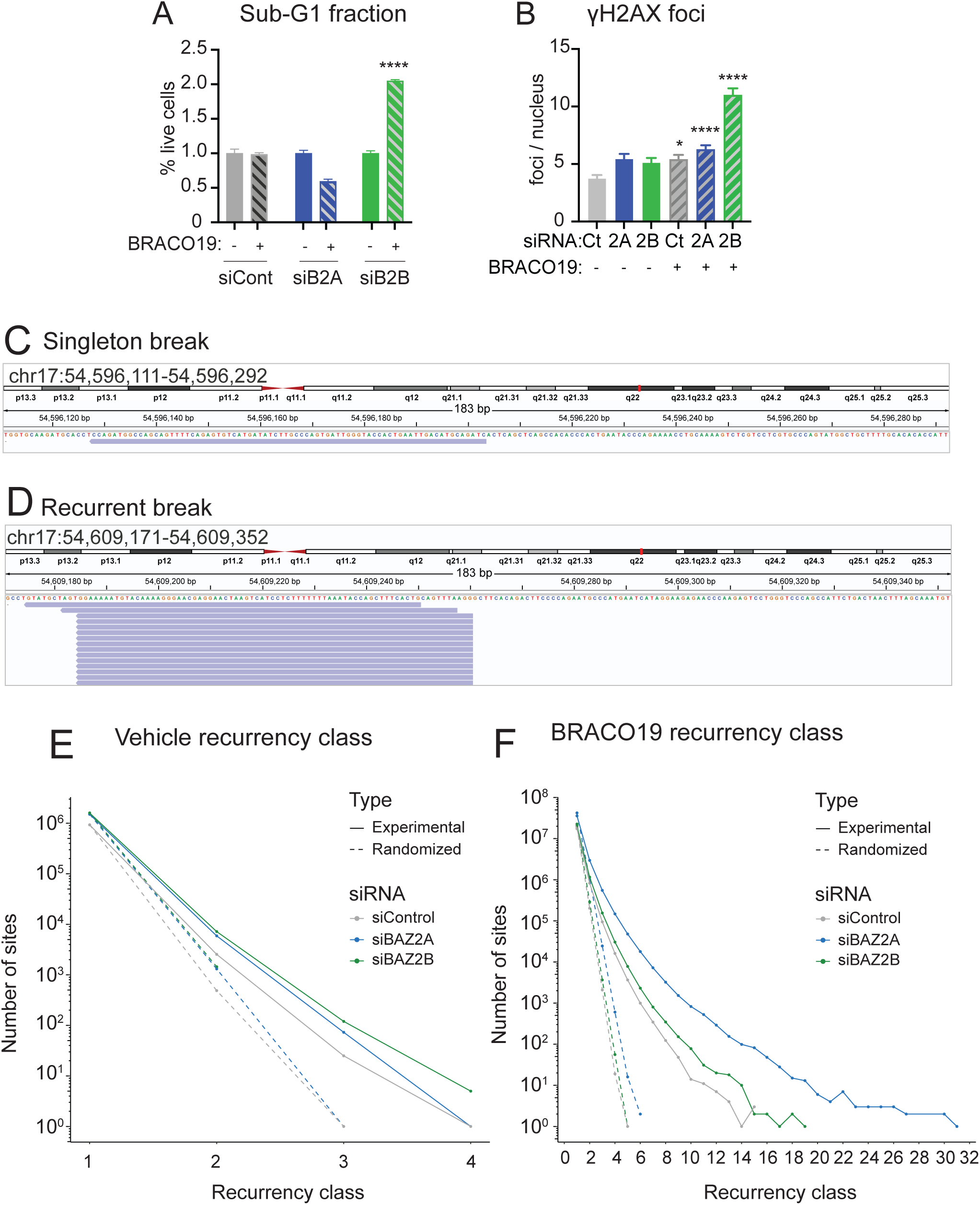
DNA Damage in the Absence of BAZ2 Chromatin Remodeling Factors. **A.** Quantitation of the sub-G1 fraction following 24 hr treatment with 10 μM BRACO19 with indicated siRNA for 72 hr. **B**. Quantitation of γH2AX foci per nucleus following 24 hr treatment with 10 μM BRACO19. **C-D**. Genome browser views showing examples of a singleton (**C**), and recurrent (**D**) double-strand breaks (DSBs) identified by INDUCE-seq. **E-F.** Recurrency class distribution of INDUCE-seq DSBs in vehicle (**E**) or BRACO19 (**F**). Recurrent DSBs are defined at 3 or more DSBs occurring at the same location. siCont, siControl; Ct, Control; 2A, BAZ2A; 2B, BAZ2B. *, p < 0.05; **, p < 0.01; ***, p < 0.001; ****, p < 0.0001 (one-way ANOVA for multiple comparisons).

To explore this in more detail, we analyzed the genomic locations of DSBs under these conditions using INDUCE-seq (63), an unbiased sequencing method that detects DNA DSBs at nucleotide-level resolution. Because INDUCE-seq is PCR-free, each read corresponds to an individual break labelled within the cell. This allows us to detect both singleton and recurrent DSBs, the latter defined as multiple DSBs occurring independently at the same location in different cells (**Fig. 4C-D**). Consistent with our IF data, BRACO19 treatment led to an increase in both singleton and recurrent DSBs across all conditions (**Supplementary Fig. 11A-B**), although not all increases achieved statistical significance. Recurrent DSBs, defined here as more than three DSBs at the same location, can be further annotated by recurrency class which denotes the number of DSBs occurring at a specific location. In the absence of BRACO19 treatment, DSB recurrency levels were not significantly different from randomized controls (**Fig. 4E**). However, BRACO19-treated samples exhibited significant increases in DSB recurrency, particularly in the highest recurrency classes, compared to the randomized controls across all conditions (siControl: *p* = 0.0048; siBAZ2A: *p* < 0.0001; siBAZ2B: *p* < 0.0002, Kolmogorov-Smirnoff test), highlighting the role of BRACO19 in genome stability (**Fig. 4F**).

### NMF Reveals BRACO19-Associated DSB Sequence Patterns

To examine the sequence context around DSBs captured by INDUCE-seq, we extracted the 5-base pair (bp) sequences surrounding each DSB and applied nonnegative matrix factorization (NMF). This represents, to our knowledge, the first application of NMF to analyze sequences surrounding DSBs, though it has been widely used to identify signatures of somatic mutations from cancer genomes (64–66). Using this approach, we identified four sequence-based components that distinctly contribute to each experimental condition (**Fig. 5A**), which resulted in samples clustering first according to BRACO19 treatment and second according to siRNA within the BRACO19-treated samples (**Supplementary Fig. 11C**). Notably, BRACO19 treatment led to an increased relative contribution of Component D and a decrease in the relative contribution of Component A across samples, irrespective of siRNA treatment (**Fig. 5A**). Of the four components, Component D had the highest GC content (54.3%) while Component A was lower (50.4%, **Fig. 5B-C**), suggesting that GC-rich sequences may contribute to the DSBs found in Component D. Examination of the 5 highest contributing 5-base pair motifs (5-mers) in each component revealed that those in Component D have some commonalities (**Fig. 5D-E**). The consensus sequence is “GT(C/G)N(C/T)” if both the sequence and the reverse complements of the top five 5-mers are considered (**Fig. 5E**), suggesting that this 5-mer makes a significant contribution to Component D and therefore BRACO19-dependent breaks.

**Figure 5:**
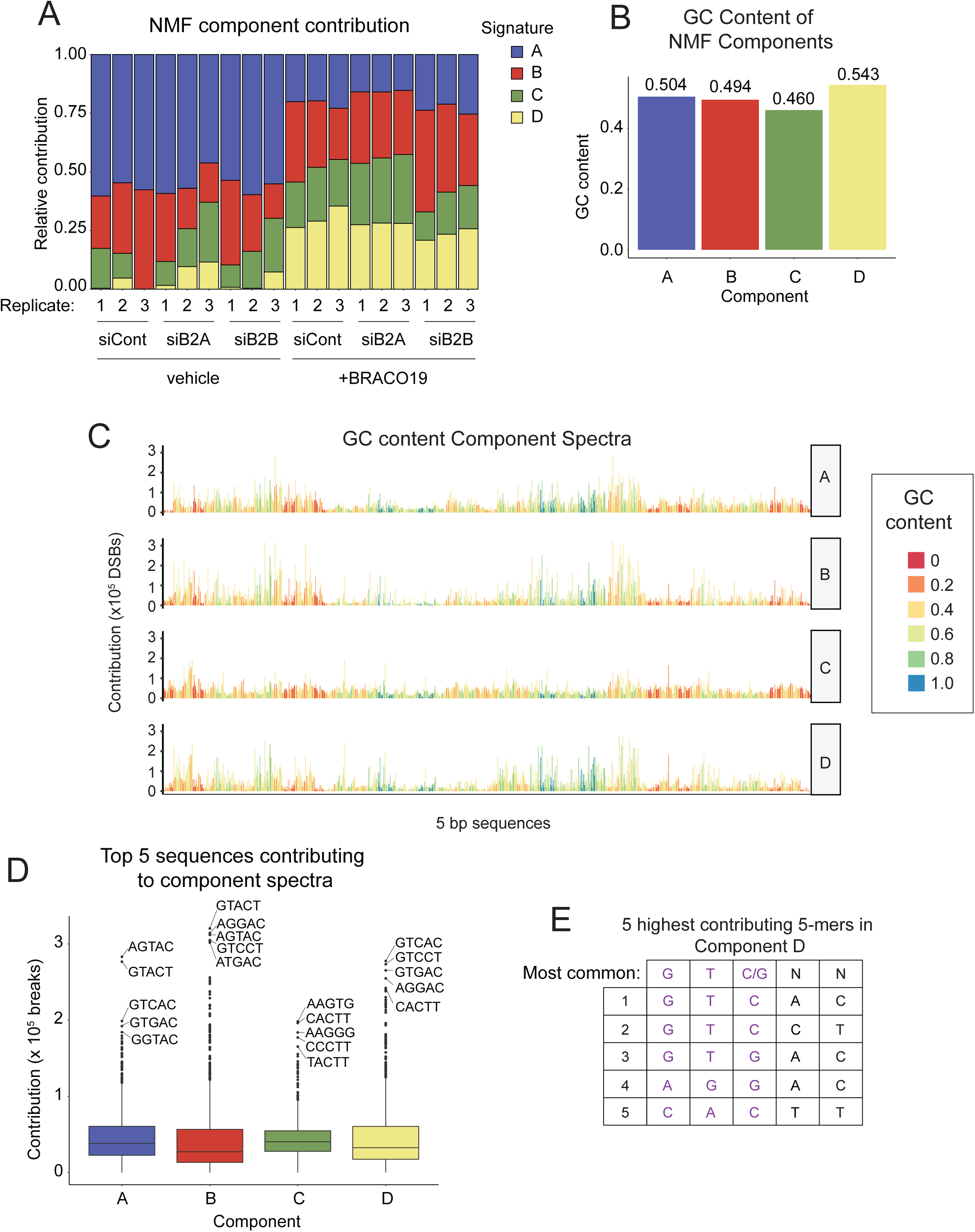
Nonnegative Matrix Factorization Reveals BRACO19-Associated DSB Sequence Patterns. **A.** Relative contribution of latent components extracted by non-negative matrix factorization (NMF) to the overall DSB pattern observed for each sample. **B.** Weighted average GC-content across NMF components. **C.** NMF component spectra, representing the contribution of each 5 bp sequence to each component; color indicates GC content. **D.** Quantitation of 5-mer sequence contributions to NMF components; the sequences of the top 5 contributing 5-mers are shown. **E**. Consensus sequence of the 5 most contributing 5-mers in Component D. siCtrl, siControl; siB2A, siBAZ2A; siB2B, siBAZ2B.

### DSBs exhibit distinct patterns around and within G4 structures

To explore the potential involvement of G4s in DSB formation, we analyzed the overlap between our INDUCE-seq and G4 CUT&Tag datasets. We detected a statistically significant enrichment of singleton DSBs at G4 peaks relative to chance by GAT analysis (*p* = 0.001, 1,000 permutations, (60)) in all siRNA conditions without BRACO19 treatment (**Fig. 6A, Supplementary Fig. 12A**). However, BRACO19 abrogated these enrichments. For recurrent DSBs (observed independently more than three times), we found no enrichment at G4 CUT&Tag peaks above random levels in the absence of BRACO19 (**Fig. 6B**). However, BRACO19 treatment resulted in a statistically significant enrichment (*p* = 0.001, GAT analysis, 1,000 permutations; **Fig. 6B, Supplementary Fig. 12A**), marking the strongest category of enrichment observed. This suggests that BRACO19 induces the most substantial association between DSBs and G4 structures in recurrent break regions.

**Figure 6:**
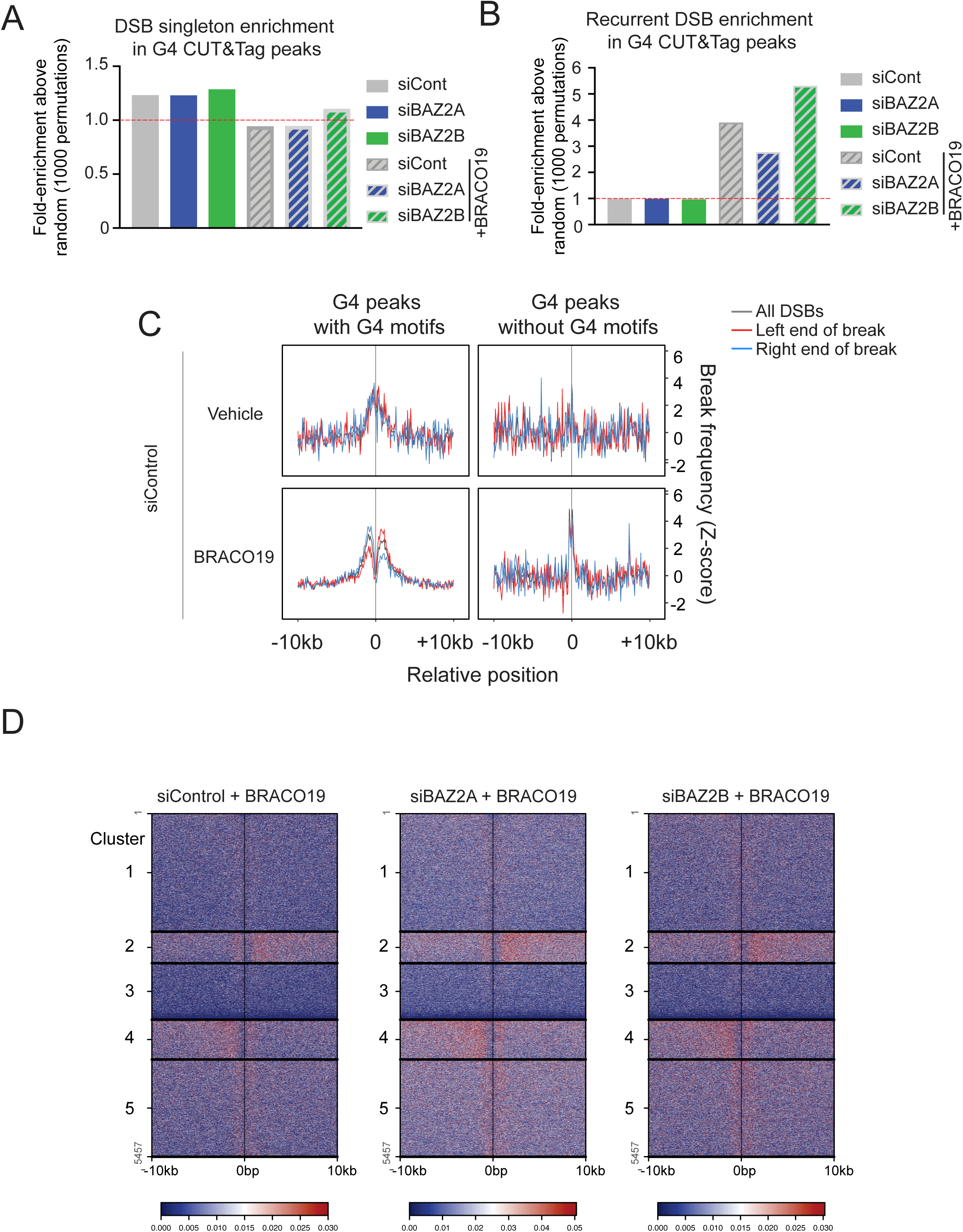
Enrichment and Distribution of DSBs in Proximity to G4 DNA Structures. **A-B.** Fold enrichment over random (GAT analysis, *n* = 1,000 permutations) for singleton DSBs (**A**) and recurrent DSBs (**B**) in G4 CUT&Tag peaks. Red line indicates a fold-change of 1 **C.** Average profile plots of DSB frequency (Z-score) within 10 kb of G4 CUT&Tag peaks with and without G4Catchall detected motifs with vehicle- or BRACO19-treatment; grey is all DSBs, red is the left end of DSBs and blue is the right end of DSBs. **D.** Unsupervised k-mean clustered heatmaps of BRACO19-associated DSBs with indicated siRNA within 10 kb of G4 peaks (all conditions combined, *n* = 5,457).

If we examine DSB frequency in a window of 20 kb surrounding the G4 structures identified by CUT&Tag, a pattern emerges. Without BRACO19 treatment, cumulative analysis of DSB frequency near G4 CUT&Tag peaks containing G4Catchall motifs displayed an elevated normalized DSB frequency, with enrichment spanning roughly 2 kb on both sides of the G4 across all siRNA treatments (**Fig. 6C, Supplementary Fig. 13A**). This enrichment was not observed at G4 CUT&Tag peaks lacking a G4 motif (**Fig. 6C, Supplementary Fig. 13A**). BRACO19 treatment led to a marked depletion of DSBs directly at the G4 site, with bimodal enrichment patterns arising on either flank (**Fig. 6C, Supplementary Fig. 13A**), giving context to our observation that BRACO19 abrogates DSB enrichment directly at the G4 peak (**Fig. 6A**). We observed a modest decrease in DSB frequency at the center of the G4 peak without BRACO19 treatment, suggesting that G4 structures induce a bimodal DSB distribution, which is intensified by BRACO19. Our analysis of BRACO19-treated samples revealed an asymmetry in DSB distribution (**Fig. 6C, Supplementary Fig. 13A**). Right ends of DSBs occurred more frequently upstream of G4 peaks, while left ends displayed the opposite trend. To assess effects of sample size, we generated subsets of the INDUCE-seq data by random down-sampling of DSBs in BRACO19-treated samples to match the smaller sizes of either the vehicle-treated samples or the G4 peaks without a G4 motif. These smaller subsets yielded similar DSB patterns to the complete datasets (**Supplementary Fig. 13B**) indicating a robust pattern that implies that G4 structures engage with or orient a directional process that drives selective DSB formation.

Closer examination of DSB distributions near G4 CUT&Tag peaks revealed a variety of distinct DSB patterns (**Supplementary Fig. 14A**). Unsupervised k-means clustering of DSB patterns relative to G4 peaks in the BRACO19-treated samples extracted five clusters corresponding to the following classes (**Fig. 6D**): 1) G4s with an equal bimodal DSB distribution flanking a pronounced trough, 2) G4s with an unequal bimodal distribution with increased DSB frequency downstream, 3) G4s with no DSBs or a very low frequency of DSBs, 4) G4s with an unequal bimodal distribution with increased DSB frequency upstream, and 5) G4s with an equal bimodal DSB distribution flanking a shallow trough. Interestingly, DSB patterns remained consistent irrespective of BAZ2 depletion, highlighting the robustness of these patterns. Additionally, the reproducible clustering of G4 peaks across samples indicates that the associated DSB patterns are intrinsic to these G4 loci. While samples from untreated cells had too few DSBs to perform unsupervised clustering, we sought to investigate whether DSB patterns induced by BRACO19 resemble those present at the same loci under endogenous conditions. Thus, we compared the DSB patterns in untreated samples at the same loci defined in the BRACO19 clusters (**Supplementary Fig. 14B**). The data revealed that, except for Clusters 1 and 5, the DSB patterns in untreated cells mirrored those observed with BRACO19 treatment, suggesting that the majority of G4-proximal DSB patterns induced by BRACO19 are also found endogenously. Notably, Cluster 1 represents one of the smallest proportions of the total population (**Supplementary Fig. 14C**). Analysis of G4Hunter scores for each cluster revealed that Cluster 3, G4s with no DSBs or a very low frequency of DSBs, had a significantly lower G4Hunter score than the other clusters (**Supplementary Fig. 15A**). Conversely, Clusters 2 and 4 which featured asymmetric patterns, had significantly higher G4Hunter scores. Together, these findings suggest that G4 structures can contribute to genome instability through a variety of mechanisms.

We hypothesized that specific genomic features might be linked to the distinct DSB patterns observed around G4s. Since a large proportion of G4 structures are found at promoters (**Supplementary Fig. 6A**), we examined DSB distributions within a 10 kb window around promoters with and without G4 structures. Promoters without G4s exhibited a strong DSB peak at the promoter and a smaller shoulder in the direction of the gene (**Fig. 7A**). By contrast, promoters containing G4s showed a secondary DSB peak upstream of the promoter, along with a more prominent shoulder extending downstream toward the gene (**Fig. 7A**). No BAZ2-dependent differences in DSB distribution around promoters were observed (**Supplementary Fig. 15B-C**). Collectively, these data suggest that the distinct pattern featuring an upstream peak and a stronger genic shoulder of DSBs, can be attributed to the presence of a G4 structure within the promoter. Since G4-containing promoters are linked to genes with higher expression levels (*p* < 0.0001, *t*-test, **Supplementary Fig. 16A**) ((67), https://depmap.org/portal), we generated a control set of genes associated with promoters lacking a G4 structure that have statistically indistinguishable mean expression levels to those of G4-containing promoters (ns, *t*-test, **Supplementary Fig. 16B**). In this expression-matched promoter set lacking a G4, we observed a peak with a shoulder (**Supplementary Fig. 16C**) that was similar to the overall G4-deficient dataset, suggesting that the observed changes are due to the presence of G4s rather than elevated expression levels. We noted that the pattern of DSBs at promoters containing a G4 (**Fig. 7A**) was similar to patterns found in Clusters 2 and 4 (**Fig. 6D, Supplementary Fig. 14B**). Analysis of defined genomic elements (58) associated with the G4s across the five clusters revealed that Clusters 2 and 4 had the highest proportions of promoters and TssAs (94.5% and 96.3% respectively). However, other clusters also showed substantial proportions of promoters and TssAs (**Supplementary Fig. 17A**), indicating that multiple distinct patterns of DSBs can be associated with gene regulatory elements.

**Figure 7:**
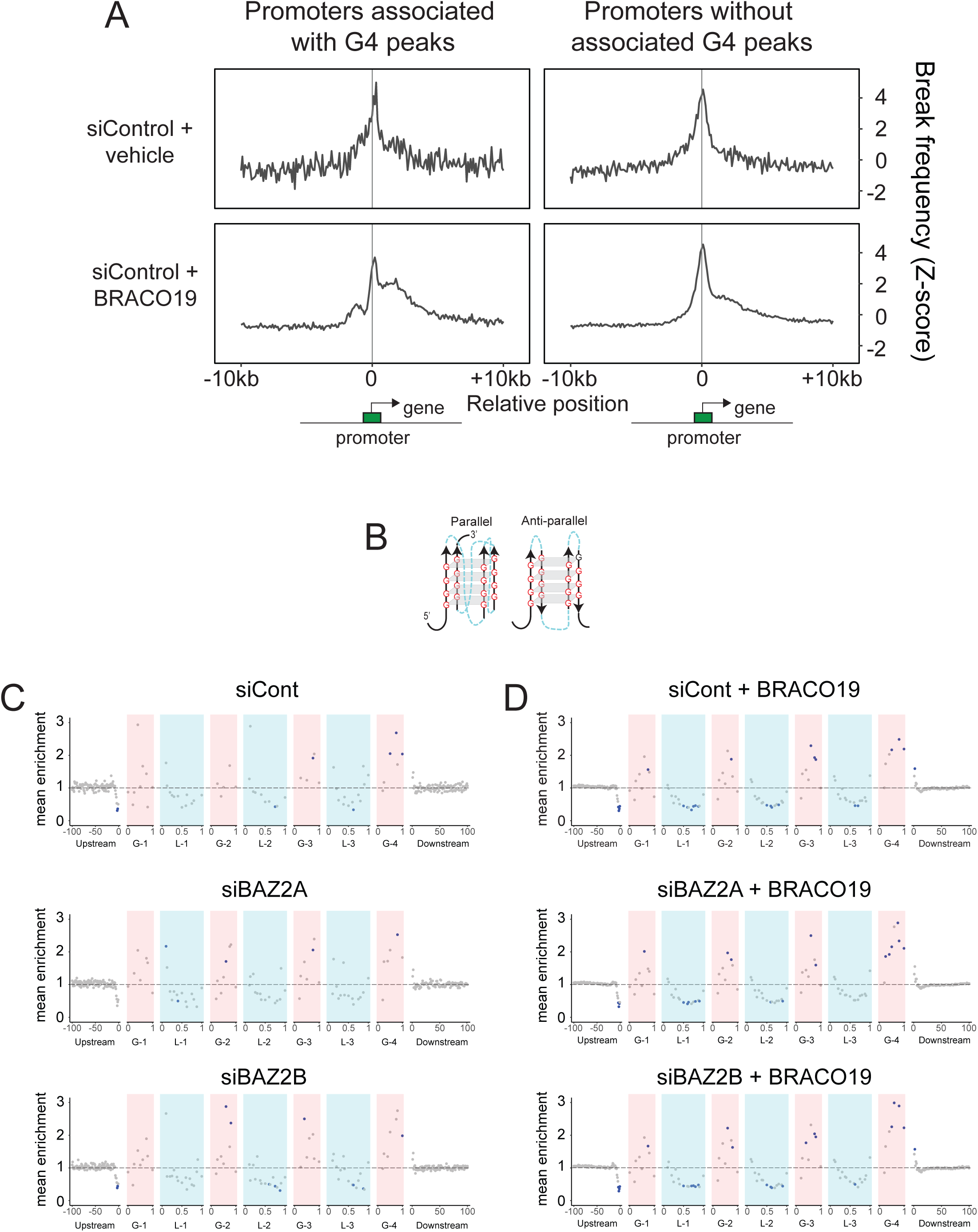
Patterns of DSBs within G4 motifs. **A.** Average profile plots of DSB frequency (Z-score) within 10 kb of promoters associated with G4 CUT&Tag peaks and promoters without associated G4 CUT&Tag peaks in vehicle- or BRACO19-treated siControl samples. **B.** Schematic representation of parallel and antiparallel G4 structures highlighting G repeats (red) and loops (blue). **C-D.** Enrichment of DSBs above random within 100 bp of G4 motifs for vehicle-(**C**) and BRACO19-treated (**D**) cells. Positions with DSB frequency significantly different relative to randomized control (fold change ≤0.5 or ≥1.5 and *p* < 0.05) are indicated in blue.

Because INDUCE-seq captures a large number of DSBs with nucleotide-level precision, we could explore patterns of DSB occurrence within G4 structures. For this analysis, however, we needed a larger G4 dataset than those found in G4 CUT&Tag peaks. We reasoned that CUT&Tag might only be detecting a subset of highly stable G4 structures, while other potentially more transient G4s could also be influencing genome stability. Therefore, we used G4Catchall (56) to identify genome-wide G4 motifs and determined all DSBs that occurred within 100 bp of these motifs. Using this expanded dataset, we mapped DSB frequency from each condition onto a length-normalized diagram of a G4 motif (**Fig. 7B-D**). We observed similar patterns in DSB frequency both with and without BRACO19 treatment (**Fig. 7C-D**). However, BRACO19-treated samples showed more statistically significant locations (**Fig. 7D**) due to the overall increase in DSBs (**Supplementary Fig. 12A**). In all conditions, DSB frequency was reduced ∼10-12 bp upstream (5’) of the G4, whereas sites immediately downstream (3’) of the G4 showed an elevated DSB frequency (**Fig. 7C-D**). The G4 loop sequences, particularly the center of each loop, had fewer DSBs than expected by chance, while the G repeats were enriched for DSBs. DSB frequency was highest in the fourth G repeat and, though still significantly elevated compared to random, lower in the other G repeats (**Fig. 7C-D**). Although no statistically significant differences in DSB patterns were linked to BAZ2 or BRACO19, we did note a trend toward DSB enrichment at the 5’ edge of loops that was reduced following BRACO19 treatment. Furthermore, BRACO19 treatment appears to enhance the upward trend in DSB frequency from G repeat 1 to G repeat 4. Taken together, these data indicate that the regions immediately upstream and within loops are depleted of DSBs, whereas G repeats and the downstream sequences are enriched for DSBs.

## Discussion

In the present study, we expanded our understanding of the regulatory relationship between chromatin modifications and G4 structure formation. Our screen identified nine chromatin modifications that suppress G4 structures, seven of which represent novel relationships and open new avenues for investigation. We characterized the BAZ2 family of chromatin remodelers, showing that their suppression of G4 structures is likely direct. Additionally, we demonstrated that aberrant G4 structures that form in the absence of BAZ2 contribute to genome instability, particularly upon stabilization by a small-molecule G4 ligand. Through in-depth analysis, we explored the genomic instability landscape induced by the G4 ligand BRACO19 and suggest that it may be a useful tool for amplifying G4-dependent mutagenesis as it allowed us to discern G4-dependent DSB patterns near promoters and within G4 structures. However, we found that BRACO19 induces both G4-dependent and -independent DSBs and introduced the use of non-negative matrix factorization (NMF) to uncover potential sequence determinants of these DSBs. Taken together, our study increases our understanding of the mechanisms governing G4 structure regulation within its chromatin context and highlights the importance of maintaining proper G4 dynamics.

We identified nine chromatin modifications that regulate G4 DNA structure formation, with several findings aligning with existing literature. Our data confirm reports that histone deacetylase inhibition increases global G4 structures (19, 21). Also, we found that inhibition of DNA methylation promotes G4 structure formation, supporting reports of an anti-correlation between G4 structures and DNA methylation (22, 68). The prevailing model suggests that G4 structures inhibit DNA methylation by sequestering DNA methyltransferase enzymes (69–71). Interestingly, our observation that DNMT1 inhibition increases G4 structures, which was corroborated by a recent study during the preparation of this paper (72), supports the notion of a reciprocal regulatory relationship in which G4 structures inhibit DNA methylation while DNA methylation constrains G4 formation. The discovery of additional chromatin modifiers like LSD1 and PRMT1, which alter histone methylation patterns, as novel regulators of G4 structures represent interesting avenues for future research.

Although G4 structures form independently of transcription (1, 73), they are important for transcriptional regulation and are linked to highly expressed genes (1, 19). ISWI complexes, which include BAZ2, regulate transcription (45, 74), and our findings indicate that BAZ2 may achieve this in part by remodeling G4 structures. BAZ2 activity could disrupt G4s or hinder their interactions with transcription factors (75). Despite their transcriptional roles, G4s contribute to genomic instability, and the absence or inhibition of BAZ2 may result in persistent G4s that promote abnormal transcription or genome instability. This connection may offer insights into the potential contribution of BAZ2B haploinsufficiency and hypermethylation to neurodevelopmental disorders including developmental delay and autism spectrum disorder (76–78). Conversely, BAZ2A over-expression has been linked to multiple malignancies including prostate and liver cancer (42, 79). Finally, BAZ2 control of G4 structures may have implications for the use of small molecule inhibitors like GSK2801 in combination oncology therapies (80, 81) or in regenerative medicine (82).

Our data indicate that loci where BAZ2B localizes are significantly enriched for G4 structures when BAZ2B is absent. This observation suggests that BAZ2B could play a role in directly suppressing G4 formation and might even be recruited to chromatin through a G4 interaction. *In vitro* and proteomic studies lend credibility to this hypothesis. For example, SMARCA5, a BAZ2 partner (39), has been shown to interact biochemically with the Pu27 G4 in the c-Myc promoter (40). Consistently, quantitative proteomics revealed that SMARCA5 interacts with the c-Kit promoter G4 structure in human cells (83). Similarly, Zhang *et al.* applied a functionalized G4 ligand for proximity labeling and found that BAZ2A interacts with endogenous G4 structures, supporting the idea that G4s might serve as docking sites for BAZ2 family complexes (84). The possibility that the BAZ2/SMARCA5 remodeling complex is recruited to or retained on chromatin by direct interaction with G4 structures merits further investigation. A key biochemical activity of BAZ2 remodeling complexes is to slide nucleosomes into nucleosome-deficient regions (39) such as G4 structures. Therefore, it is plausible that BAZ2 complexes are recruited to G4-rich areas, where they might reposition nucleosomes to suppress G4 formation. However, not all G4-containing genes are bound by BAZ2B (**Fig. 3F**) indicating that additional, yet unidentified, signals are required for its recruitment.

In addition to recruiting proteins, G4 structures can also induce genome instability and our data sheds light on patterns of G4-dependent DSBs. We find that G4-containing promoters exhibit higher DSB frequencies upstream and downstream compared to those without G4s (**Fig. 7A**). The DSB distribution that we observe in promoters lacking G4 structures resembles the DSB patterns reported in recent studies of DSB distribution around transcription start sites in mouse and human cells (85, 86), suggesting that the sharp peak and the downstream shoulder of DSBs toward the gene are universal features of transcription hubs. By intensifying the downstream shoulder of DSBs, BRACO19 may amplify an intrinsic promoter-associated pattern of DSBs. Based on similar patterns of topoisomerase (TOP2A and TOP2B) occupancy, Murat *et al.* suggest that the DSB pattern that they observe could be mediated by topoisomerases (85). Interestingly, BRACO19 uniquely induces an upstream DSB peak in G4-structured promoters which could correspond to topoisomerase activity aimed at restoring negative supercoiling. Topoisomerase I (TOP1), which can alleviate negative supercoils (87), interacts with and is inhibited by G4 structures (88, 89). BRACO19 might disrupt TOP1 binding to G4 structures, allowing more TOP1 activity around G4-enriched promoters. Consistent with a role for topoisomerases in G4-mediated DSBs, a recent screen identified TOP2A as a major effector of the cytotoxicity produced by the G4 ligands PDS and CX-5461 (90). Mechanistically, BRACO19-induced stabilization of G4 structures could amplify stress generated during transcription or liberate topoisomerases, triggering an increase in local topoisomerase activity.

Dissecting the molecular processes that contribute to DSB formation at G4s is complicated by the finding that G4s are hubs for several cellular processes beyond transcription. For example, G4 structures have been proposed to coincide with human replication origins (5), although consensus on this association is lacking (91). Interestingly, we find an asymmetric pattern of DSBs around G4 peaks (**Fig. 6C**), with right ends of DSBs predominating upstream (5’) and left ends of DSBs being more common downstream (3’). Since replication fork collapse produces one-ended DSBs, this asymmetry may reflect fork collapse near G4-associated origins, where upstream forks collapse into right ends of DSBs, and downstream forks produce left ends of DSBs (**Supplementary Fig. 17B**). If this specific pattern of DSBs is replication-dependent, it might suggest that G4 stabilization by BRACO19 influences origin usage or increases torsional stress at replication origins inducing more collapsed forks. Further experimental investigation is needed to clarify this relationship.

We found that within G4 motifs, G repeats were enriched in DSBs, whereas loop sequences were relatively depleted of DSBs (**Fig. 7C-D**). In contrast, a recent study using human-orangutan divergence and human substitution frequencies found both loops and G repeats exhibited substitution rates above baseline, with loops displaying a higher rate than G repeats (92). One possibility for this inverse pattern of susceptibility is that different regions within the G4 structure may be predisposed to distinct types of mutational events or the mitigating DNA repair processes. An alternative explanation is that selective pressures differ between G repeats and loops, with loops potentially accommodating substitutions without compromising G4 formation, whereas G repeats are crucial for G4 formation. Consequently, the patterns reported by Guiblet *et al.* could reflect different selective pressures on G4 substructures. Our observation that DSB frequency is highest at the 3’ end of the G4 structure may reflect stalling of an enzyme such as a helicase at the 3’ side of the structure. Perhaps, once it is past this 3’ region, it can unwind the rest of the G4 more efficiently.

This study identifies novel chromatin modifications that regulate G4 DNA structures and highlights BAZ2 complexes as direct G4 suppressors. The unique patterns of genomic instability, amplified by BRACO19, offer insights into G4-dependent DSBs and open avenues for further exploration of G4-dependent mutagenesis.

## Data availability

The CUT&Tag and INDUCE-seq datasets generated in this study will be deposited in the NCBI GEO database under an accession number that will be made publicly available upon publication. Publicly available datasets used in this study include the OQ data generated in the presence of PDS (downloaded in BED format from GEA, accession GSM3003540), and BAZ2B mouse ChIP-seq data (downloaded in FASTQ format from GEO, accession GSM5558422). The hg19 and dm6 reference genomes were retrieved from the UCSC database. The hg19 blacklist file (identifier ENCFF200UUD) was obtained from the ENCODE database. Genome annotation for A549 cells (file E114_25_imputed12marks_dense.bed.gz) was retrieved from the Roadmap Epigenomics database.

## Code availability

Custom scripts developed in Python, R, and Bash for this study will be made publicly available on GitHub upon publication. The G4Catchall script was obtained from https://github.com/odoluca/G4Catchall, the G4-Cluster code from https://github.com/UofLBioinformatics/G4-Cluster, and the GAT analysis code from https://github.com/AndreasHeger/gat.

## Acknowledgements

We thank the members of the Day laboratory for helpful discussions. This work was supported by an NSF CAREER Award 2143016 (T.A.D.). We thank the Institute for Chemical Imaging of Living Systems at Northeastern University (CILS) for microscope support and maintenance.

## Materials and Methods

### Cell culture and reagents

A549 cells (ATCC, CCL-185) were cultured in DMEM (Corning, 10-013-CV) supplemented with 10% fetal bovine serum (FBS; Gibco, 10437028), 100 U/mL penicillin and 100 μg/mL streptomycin (Gibco, 15140163). All cells were cultured in a humidified incubator with 5% CO2 at 37°C. Cells were treated with inhibitors listed in **Supplementary Table 1** at the indicated concentrations (HY-L005, MedChem Express).

### Construction of HA-BAZ2B cell line

To create a homozygous HA-tagged A549 cell line, 2 × 10^5^ cells were nucleofected (Lonza, V4XC-2032) with 75 pmol crRNA (**Supplementary Table 2**), 75 pmol tracrRNA (IDT, 1072532), 125 pmol Cas9 enzyme (IDT, 1081058) and 4 μM HDR Donor Oligo containing the template for the modified HA-tag including a NdeI restriction site within the tag (**Supplementary Table 2**). Cells were then grown in four wells of a 96-well plate for 24 hours in media containing HDR enhancer V2 (IDT, 10007910). Single-cell clones were isolated and screened for insertion of the HA tag by PCR and restriction digestion with NdeI (NEB, R0111S). The insertion was also verified by nanopore sequencing of the unrestricted PCR amplicon.

### siRNA transfection

A549 cells were transfected with 71-86 nM siRNA using lipofectamine 3,000 transfection reagent (Fisher Scientific, L3000015) for 48-72 hours. The siRNA sequences are listed in **Supplementary Table 2**.

### BG4 production

The BG4 production was adapted from the protocol described by De Magis *et al.* (93). Briefly, the pSANG10-3F-BG4 plasmid, a gift from Shankar Balasubramanian (Addgene, #55756; http://n2t.net/addgene:55756; RRID: Addgene_55756), was transformed in BL21(DE3) competent cells (Thermo Scientific, EC0114). Transformed cells were grown in 2 L YT media (Sigma-Aldrich, Y2377-250G) with 50 μg/mL Kanamycin (Fisher Scientific, BP906-5) at 37°C at 220 RPM for approximately 9 hours. The expression of the BG4 antibody was then induced by adding 0.5 mM Isopropyl-β-D-thiogalactopyranoside (MP Biomedicals, 114064112) and incubating overnight at 25°C. Bacteria were harvested by centrifugation at 4,500 x *g* at 4°C for 12 min and lysed in TES buffer (50 mM Tris-Cl pH 8.0, 1 mM EDTA, 20% sucrose) on ice for 10 min. The lysate was then spun at 16,000 x *g* at 4°C for 1 hr and the supernatant was filtered through a 0.45 μm vacuum filter. BG4 was then purified on a Ni-NTA Sepharose column (Thermo Fisher, 88226) pre-equilibrated with TES buffer. The column was then washed twice with PBS supplemented with 100 nM NaCl and 10 mM Imidazole (pH 8.0) by centrifugation at 700 x *g* at 4°C for 2 min. BG4 was eluted in PBS with 250 mM Imidazole and EDTA-free cOmplete Protease Inhibitor Cocktail (Roche, 11873580001). The elution buffer was then exchanged by inner cell salt buffer (25 mM HEPES pH 7.6, 110 mM KCl, 10.5 mM NaCl, 1 mM MgCl2 through overnight dialysis (Thermo Fisher, 66830) whereafter the BG4 was concentrated using the Amicon Ultra-15 centrifugal Filter unit with 10 kDa cutoff (Millipore Sigma, UFC901024). The purity and concentration of the BG4 preparation was determined by SDS-PAGE.

### Real-time RT-qPCR

Total RNA was isolated using the Monarch Total RNA Miniprep Kit (NEB, T2010S). The qRT-PCR assay was performed using 500 ng of total RNA and the Luna Universal One-Step RT-qPCR Kit (NEB, E3005S) on a QuantStudio 3 system (Applied Biosystems). The qRT-PCR conditions were 55°C for 10 min, 95°C for 1 min, 45 cycles of 95°C for 10 sec and 60°C for 1 min. The mRNA was quantified using the comparative CT Method normalized to ACTB levels. The primers are listed in **Supplementary Table 2**.

### Western Blotting

Cells were harvested and lysed with RIPA buffer (Thermo Fisher, 89901) containing protease inhibitor (Thermo Fisher, 78430) for 5 min on ice. They were subsequently sonicated for 4 cycles (30 sec on, 30 sec off) using the Bioruptor Pico (Diagenode) and centrifuged for 15 min at 14,000 x *g*. Equal amounts of protein were loaded on an 8% polyacrylamide gel and resolved by SDS-PAGE. The proteins were then transferred to a nitrocellulose membrane (Bio-Rad, 1620112) using a semi-dry transfer apparatus (Bio-Rad) 1 hr 30 min at 16 V and overnight at 5 V. Blots were blocked using 5% milk in PBST (0.05% Tween-20) for 10 min, stained with BRD9 Ab in 5% milk in PBST (1:2000; Bethyl, A303-781A) for 1 hr 30 min followed by 5 washes with PBST and staining with Goat-anti-rabbit IgG StarBright Blue 700 (1:2,500; Bio-Rad, 12004161) for 1 hr. After 5 more washes with PBST the blot was imaged using the ChemiDoc Gel Imager (Bio-Rad) and reprobed using Anti-Actin hFAB Rhodamine Ab (1:2,500; Bio-Rad, 12004163). Band intensities were quantified using the FIJI (94) software and the BRD9 levels were normalized to the Actin signal.

### Immunofluorescence

At the indicated times after inhibitor treatments or transfections, cells were washed twice with PBS. For the initial BG4 screening one pre-extraction step was included with CSK buffer supplemented with protease inhibitor and 1 mM ATP for 10 min on ice. The cells were then fixed with 4% paraformaldehyde (Fisher Scientific, 50-980-488) for 10-15 min at room temperature. Subsequently, they were washed three times with PBS and permeabilized with 0.1% Triton X-100 in PBS for 10 min at room temperature. Where specified, an RNAse A treatment was included at a concentration of 0.24 mg/mL in PBS for 1 hr at 37°C. Cells were then blocked using 3% bovine serum albumin (BSA) in PBST for 30 min at room temperature or 0.5% goat serum in PBST for 1 hr at 37°C, followed by an incubation with 1:500 BG4 (Absolute Antibody, Ab00174-1.1) overnight at 4°C for the screen, 1:100 BG4 (EDM Millipore, MABE917; or homemade BG4) for 1 hr at 37°C, 1:1,000 γH2AX Ab (EDM Millipore, JBW301) or 1:500 HA-tag CUTANA CUT&RUN Ab (EpiCypher, 13-2010) for 2 hr at room temperature. This was followed by 3 washes with PBST (0.1% Tween-20). For the IF using BG4 (MABE917 or homemade BG4) an additional step was added to incubate the cells with 1:800 Anti-Flag Ab (Cell Signaling Technology, 2368S) followed by three washes with PBST. The cells were then incubated with 1:1,000 Alexa Fluor 594 Goat-anti-Rabbit (Thermo Fisher, A-11012) or Alexa Fluor 488 Goat-anti-Mouse antibody (Thermo Fisher, A-11001) in blocking buffer for 1 hr at room temperature. Followed by four washes using PBST and mounting using DAPI Vectashield (Fisher Scientific, NC1695563). They were imaged at a 40x magnification on a Zeiss widefield (for the initial screen), Zeiss LSM710 or LSM880 confocal microscope.

### Propidium iodide (PI) and BrdU cell cycle assay

GSK2801 treated cells were pulsed with 10 μM BrdU for 1 hr, then fixed with 35% ethanol in DMEM for 1 hr at 4°C. Subsequently, they were incubated for 20 min at room temperature in 1 mL 2N HCl, washed in 0.1 M sodium borate (pH 8.5), and washed in PBS. The cells were then stained with anti-BrdU FITC-conjugated antibody (BD Bioscience, 347583) in 0.5% Tween-20 and 0.5% BSA for 30 min at room temperature followed by staining in 0.5 mL PBS containing 8 μg/mL RNase and 50 μg/mL propidium iodide for 30 min at room temperature. The siRNA treated cells were fixed with 35% ethanol in DMEM for 30 min at 4°C, then pelleted at 1,000 x *g* for 1 min, washed once with PBS and resuspended in 0.5 mL PBS containing 8 μg/mL RNase A and 50 μg/mL propidium iodide followed by a 30 min incubation at room temperature. All flow cytometry experiments were run on the Attune Nxt Acoustic Focusing Cytometer and analyzed using Attune Cytometric Software.

### CUT&Tag

The CUT&tag protocol was adapted from Lyu et al., 2022 (16) with modifications. Briefly, 5 × 10^5^ (G4 CUT&Tag) or 1 × 10^6^ (HA CUT&Tag) cells were harvested per condition, washed with PBS and resuspended in NP40-digitonin buffer (20 mM HEPES, pH 7.5, 150 mM NaCl, 0.5 mM Spermidine, 0.05% digitonin, 0.01% NP40 and Roche cOmplete Protease Inhibitor EDTA-free). The cells were then centrifuged at 600 x *g* for 3 min, resuspended in 200 μL antibody buffer (NP40-digitonin buffer with 1% BSA, for the BG4 step only, 2 mM EDTA is added) containing 2 μg of BG4 (purified from *E. coli*), anti-H3K4me3 Ab (G4 CUT&Tag: EDM Millipore, MC315; HA CUT&Tag: EpiCypher, 13-0060) or anti-HA Ab (EpiCypher, 13-2010) and 35 ng Drosophila Spike-in Chromatin (Active Motif, 53083) and incubated overnight at 4°C. Nuclei were washed once with NP40-digitonin buffer followed by centrifugation at 600 x *g* for 4 min, whereafter the nuclei were resuspended in 200 μL antibody buffer containing 1:50 anti-Flag Ab (Sigma-Aldrich, F1804-200UG) for the BG4 samples or 2 ug anti-Rabbit Ab (G4 CUT&Tag: Antibodies Online, ABIN101961; HA CUT&Tag: EpiCypher, 13-0047) and incubated for 1 hr at room temperature with end-over-end rotation. For the BG4 samples an additional antibody incubation followed three washes by resuspending the nuclei in 200 μL antibody buffer with 1:50 anti-mouse Ab (Sigma-Aldrich, M7023) and incubating for 1 hr at room temperature. The nuclei were then washed three times and resuspended in 200 μL dig-300 wash buffer (20 mM HEPES, pH 7.5, 300 mM NaCl, 0.5 mM Spermidine, 0.01% digitonin, 0.01% NP40 and Roche cOmplete Protease Inhibitor EDTA-free) with 2.5 μL pA-Tn5 (EpiCypher, 15-1017) and incubated at room. After three washes with dig-300 wash buffer with centrifugation at 300 x *g* for 6 min, the nuclei were resuspended in 200 μL tagmentation buffer (dig-300 wash buffer containing 10 mM MgCl2) and incubated at 37°C for 1 hr. The reaction was then stopped by adding 6.7 μL of 0.5 M EDTA, 2 μL of 10% SDS and 5 μL of 20 mg/mL proteinase K and incubating at 63°C for 1 hr. Genomic DNA was purified using a DNA Clean & Concentrator-5 kit (Zymo Research, D4014), eluted in 20 μL, and RNA was digested by adding 1 μL RNase cocktail (Fisher Scientific, AM2288) with incubation for 20 min at 37°C. For the library prep, 2 μL of 10 μM uniquely barcoded universal i5 and i7 primers (**Supplementary Table 2**) as well as 25 μL NEBNext Ultra II Q5 2x PCR Master mix (NEB, M0544L) were added to the 21 μL DNA and a short PCR was performed (72°C for 5 min, 98°C for 30 sec, 13 x (98°C for 10 sec, 63°C for 30 sec), 72°C for 1 min). The library PCR products were cleaned up with Agencourt AMPure XP beads (Beckman Coulter, A63881) and sequenced at the Molecular Biology Core Facilities of the Dana-Farber Cancer institute on a NovaSeq sequencer with depths of 10-20 million reads per sample and 150 bp Paired-End sequencing.

### INDUCE-seq

Following the indicated transfections and optional 18 hr BRACO19 treatment, cells were harvested, resuspended in PBS, and plated at a density of 2 × 10^5^ cells per well onto a poly-D-lysine 96-well plate. The plates were spun at 300 x *g* followed by incubation for 2 hr at 37°C and 5% CO2. Cells were then fixed by adding methanol-free formaldehyde at a 4% concentration and incubating for 10 min. After washing twice with PBS, the cells were stored in 200 μL PBS at 4°C until the INDUCE-seq was performed as previously described (95). Briefly, cells were permeabilized with Buffer 1 (1hr 4°C; 10 mM Tris-HCl, 10 mM NaCl, 1 mM EDTA, 0.2% Triton x-100) and Buffer 2 (1hr 37°C; 10 mM Tri-HCl, 150 mM NaCl, 1 mM EDTA, 0.3% SDS) with intervening PBS washes. Cells were washed with Cutsmart^TM^ buffer (NEB, B6004S) then DNA breaks were end-repaired (1 hr room temperature; Quick Blunting Kit (NEB, E1201L) plus 100 µg/mL BSA) and A-tailed (30 min 37°C; NEBNext^®^ dA-Tailing Module; NEB, E6053L) with intervening Cutsmart^TM^ washes. A modified p5 adapter was ligated onto A-tailed break-ends in-situ using T4 DNA ligase (16-20 hr 16°C; NEB, M0202M). Excess adapter was removed with 10 washes with wash buffer (10 mM Tris-HCl, 2 M NaCl, 2 mM EDTA, 0.5% Triton x-100), then cells were lysed in DNA extraction buffer (10 min, 37°C; 10 mM Tris-HCl, 100 mM NaCl, 50 mM EDTA, 1% SDS with addition of 1 mg/ml Proteinase K) and crosslinking was reversed (1 hr, 65°C). Genomic DNA was extracted and purified (ZR-96 Genomic Clean & Concentrator-5, Zymo), then sheared to a 300-500 bp (Covaris ME220) and size selected using CleanNGS beads (GC Biotech, CNGS-0050). DNA was end-repaired and A-tailed (NEBNext Ultra II End Repair/dA-Tailing Module, NEB E7546L) and a modified p7 adapter was ligated (NEBNext Ultra II Ligation Module, NEB E7595L) to repaired break-ends. Libraries were purified twice, and size selected using CleanNGS beads (GC Biotech, CNGS-0050). Purified libraries were quantified using qPCR (KAPA Library Quantification Kit for Illumina^®^ Platforms (Roche, 07960255001) and 1 x 75 bp sequencing performed using a high output flow-cell on a NextSeq 550 (Illumina).

### Data analysis

#### CUT&Tag

Following a quality control check using FASTQC (v.0.11.9) (96), raw FASTQ reads were aligned to the hg19 and the dm6 reference genomes (downloaded from the UCSC database) using Bowtie2 (v.2.4.5) (97), the mosaic adapter sequences were trimmed during this step using the −5 19 parameters. The SAM files were deduplicated using Picard MarkDuplicates (v.2.27.4) followed by conversion to BAM format using samtools (v.1.15.1) (98) and blacklist removal from the BAM files with bedtools intersect (v.2.30.0) (99) using the ENCODE blacklist bed file for hg19 (100). Genome coverage tracks were generated using deeptools bamCoverage (v.3.5.1) (101) with the following parameters: --binSize 5 --normalizeUsing RPGC --effectiveGenomeSize 2,864,785,220. Peaks were called using MACS2 (v.2.1.4) (102) and high quality peaks were identified as peaks present in two or more replicates generated with bedtools intersect. For SEACR peak calling an additional step was performed: based on the sequencing depth of the spike-in alignment (seqDepth), a scaling factor was calculated by dividing 10,000 by the seqDepth value. The BAM files were then converted to bedgraph format using bamCoverage with the same parameters as above and --scaleFactor followed by the calculated scaling factor. The peaks were then called using SEACR (v.1.3) (103) with parameters 0.01 non stringent. Randomized size matched datasets were generated using bedtools shuffle with parameters: -seed 123 -chrom -excl blacklist_for_hg19.

#### INDUCE-seq

Sequencing reads were processed as previously described (95). Briefly, read trimming was performed using Trim Galore (https://www.bioinformatics.babraham.ac.uk/projects/trim_galore/) to remove adaptor sequences, followed by alignment to the human genome (hg19) using BWA-mem (104). Soft-clipped reads, low-quality alignments (MAPQ < 30), secondary alignments, optical duplicates and alignments mapping to the ENCODE blacklist or incomplete reference contigs were removed using samclip (https://github.com/tseemann/samclip), samtools (98), bedtools (99), and custom awk scripts. Break positions were assigned as the first 5’ nucleotide upstream of each read with respect to strand orientation to generate a BED file of every break at a single-nucleotide position. Bedtools merge was then utilized to generate a BED file of quantified single nucleotide break positions. Recurrent breaks were identified by using bedtools merge with parameter d0 and by subsequently filtering the data for locations containing more than three breaks.

#### Peak annotation

To annotate the peaks, our peaks were overlapped with the 25-state imputation based on chromatin state model for A549 cells (E114), available through the Roadmap Epigenomics Project (105), using bedtools intersect. The distribution of our peaks across the different states was then calculated. To evaluate peak strength, we plotted the score outputted by the MACS2 in the 5th column of the narrowPeak files.

#### G4 motif analysis

Our peaks were transformed into FASTA files using the bedtools getfasta command, and the sequences were searched for recurrent motifs using MEME suite (106) with the zero or one occurrence per sequence (zoops) option. The FASTA sequences were also used to identify G4 motifs using G4Catchall (56) using the following parameters: --G2L 1..12 --max_imperfect_Gtracts 0 --G4H. The absolute values of the G4Hunter scores outputted by G4Catchall were plotted for the different conditions using the ggplot2 (107) R package. The G-run length was calculated for the first four G-runs detected in each motif (5’ to 3’ orientation) and the loop length and loop content was calculated for the first three loops in each motif. Of note, many G4 motifs had non-symmetrical G-runs (meaning the length of the G-runs were not equal within one motif). This implies that some of the guanines counted as part of the G-runs would most likely be at the start or end of a loop instead of within the G-run itself. This skews the analysis, especially when looking at the loop content, as the G% is likely artificially low because of this. Based on the chromosomal coordinates of genome-wide G4 motifs identified by G4Catchall and DSB locations, we calculated the distance between DSBs and G4s as well as the position of DSBs within G4s. The frequency of DSBs occurring within 100 bp of a G4 and at normalized locations within a G4 was calculated and visually represented as enrichment compared to a randomized control set. To obtain the normalized locations within a G4, the position of the break within the feature was divided by the length of the feature so that features of different lengths could be plotted on the same graph. G4 motifs with more than 4 G-runs were excluded for the creation of these frequency plots. To evaluate enrichment of overlap between our dataset and a previously published dataset of observed quadruplexes (OQ) following PDS treatment (108), we used the bedtools intersect tool.

#### Pathway analysis

Pathway analysis on the HA-BAZ2B peaks overlapping with siBAZ2B G4 CUT&Tag peaks was done using the ChIPseeker R package (v.1.40.0) (109).

#### Clustering

G4 clustering analysis was performed using the G4-Cluster tool published by Neupane *et al.* (110).

#### Statistical enrichment in overlap

To assess our data for enrichment in overlap, the GAT analysis (111) was performed using 1,000 permutations.

#### Non-negative matrix factorization of the sequence context of breaks

The positional information of the break data was used to retrieve the 5 bp sequence around each break site, utilizing bedtools slop and getfasta. These data were used to generate a matrix detailing the 5 bp sequence context of breaks across experimental samples. Matrix factorization was performed using the “nmf” package in R (112). The matrix was formatted and pre-processed prior to analysis. The counts were normalized to the genome-wide frequency of each 5 bp sequence. To determine the factorization rank, a range of ranks (2–10) were assessed, with 200 iterations performed for each rank, using the ‘brunet’ algorithm. The optimal rank was determined by evaluating the cophenetic correlation coefficient and residual sum of squares for each rank. NMF was then performed using this factorization rank to decompose the input matrix into two lower-rank matrices: a basis matrix representing the features of the extracted components, and a coefficient matrix representing the contribution of these components across samples.

#### Data visualization

Genome browser views were created using IGV (v.2.12.3), and the scaled Venn diagrams were created using meta-chart (https://www.meta-chart.com/). Traces showing TSS and TTS were created using the ChIPseeker R package (v.1.40.0) (109), and the heatmaps and associated z-score normalized traces were created using the SeqPlots software (v.1.12.1) (113). Coverage tracks of breaks were generated using bedtools genomecov to produce bedGraph files in variableStep format, followed by conversion to bigWig format using UCSC tools (114). The break data was loaded into seqplots as bigWig signal tracks, while G4 peak or motif positions were loaded as BED format feature files. Average profile plots and related heatmaps were generated using: 100 bp track binning, mean statistic, midpoint features, 10 kb upstream and downstream. For the profile plots, the data was z-score transformed to allow comparison between different samples.

For the heatmaps, k-means clustering was performed in SeqPlots, with cluster information obtained as a CSV. The G4 peak information relating to each cluster was extracted and formatted as a BED file. These BED files of the G4 peaks per cluster were then loaded into SeqPlots as a feature file to generate average profile plots of breaks relative to these different classes of G4 peaks.

The promoters used for the z-score normalized traces were identified using the chromatin state model for A549 cells (E114) available through the Roadmap Epigenomics Project (105), strandedness was assigned to the promoter based on the location of the nearest gene start site. The traces were then generated with the “strand on” option to account for transcriptional direction. Expression levels of the genes influenced by the nearest promoter datasets were assessed using DepMap data specific to A549 cells (115). The list of genes with matching expression levels was then extracted, and custom R code was used to down-sample the promoter without associated G4 peaks dataset to match the number of promoters, the mean expression level, and the standard deviation of the promoter dataset associated with G4 peaks.

## Notes

### Competing Interest Statement

The authors have declared no competing interest.

