## Supplementary Information for "The Role of BAZ2-dependent Chromatin Remodeling in Suppressing G4 DNA Structures and Associated Genomic Instability"

Supplementary Figures S1-S17
Supplementary Tables 1-2

Supplementary Figure 1

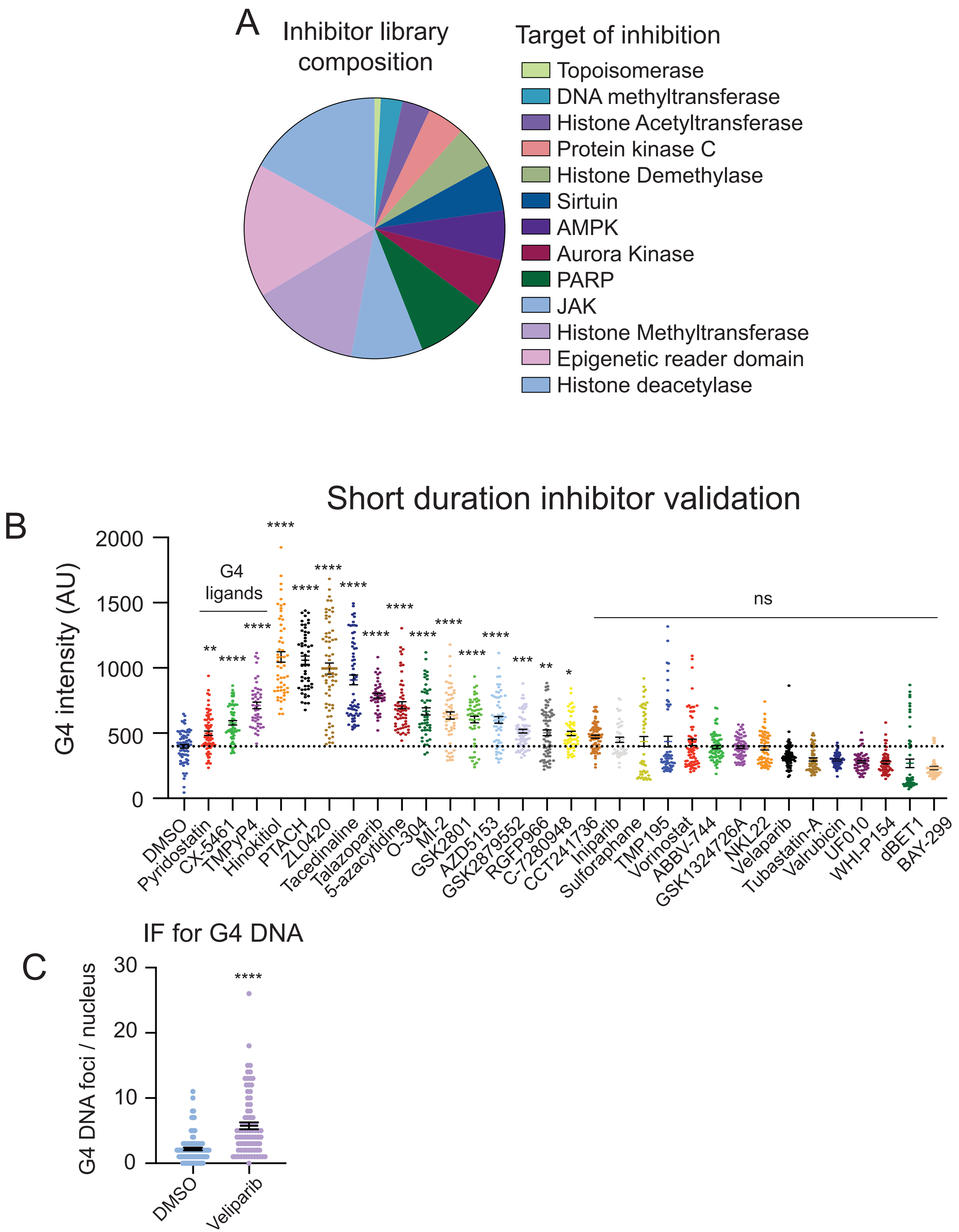

**Supplementary Figure 1: Validation of chromatin modifiers that suppress G4 DNA formation. A.** Composition of the library of small molecule inhibitors of chromatin modifying processes. AMPK, adenosine monophosphate-activated protein kinase. PARP, poly (ADP-ribose) polymerase. JAK, Janus protein tyrosine kinase. **B.** Quantitation of BG4 intensity following a 2 hr treatment with 10 μM of putative hits or G4 ligands. **C.** Quantitation of BG4 DNA foci per nucleus following a 2 hr treatment with 10 μM veliparib, one of the hits identified in the original screen, including RNAse A. ns, non-significant. *, $p < 0.05$; **, $p < 0.01$; ***, $p < 0.001$; ****, $p < 0.0001$ (One-way ANOVA for multiple comparisons).

Supplementary Figure 2

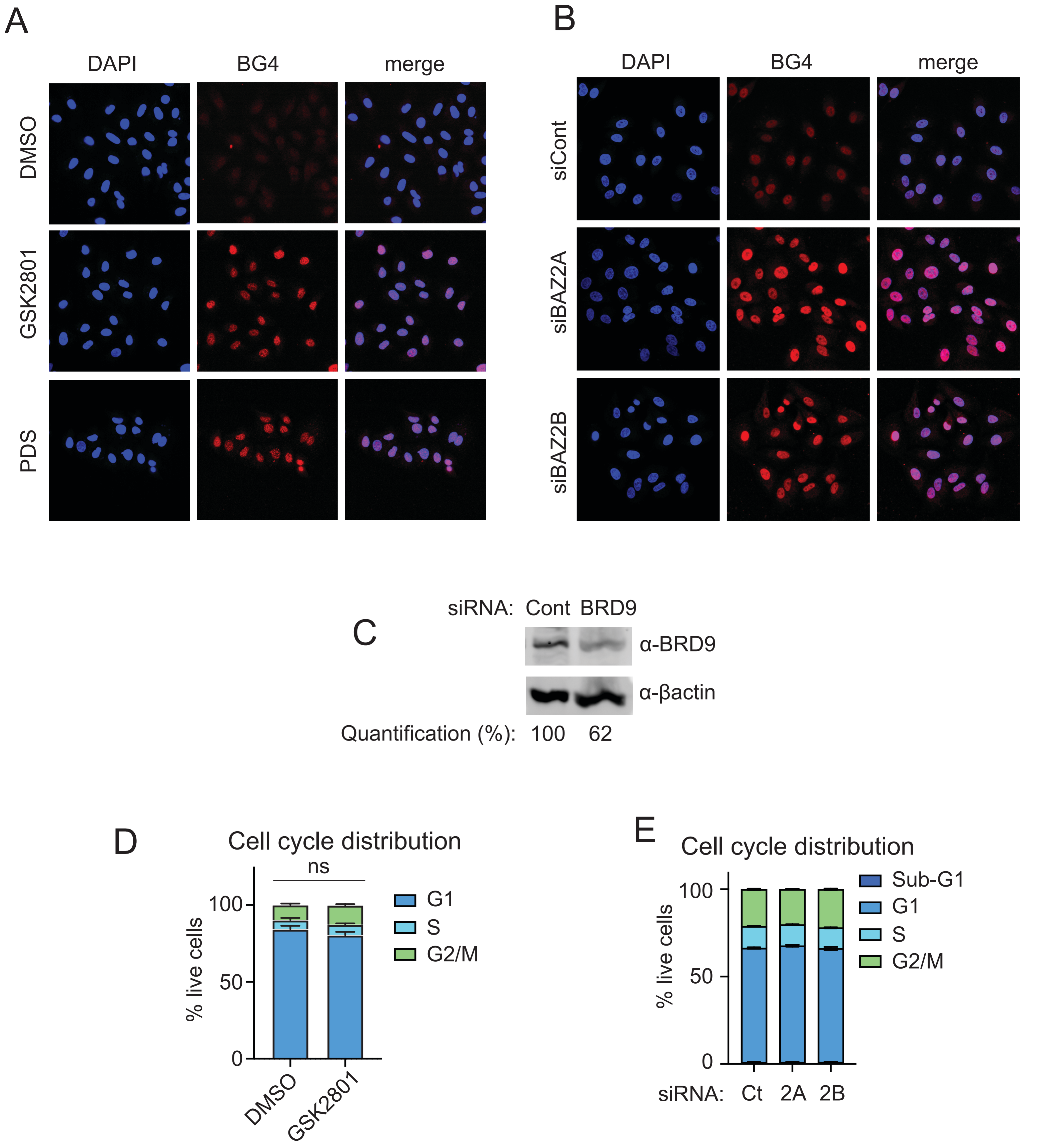

**Supplementary Figure 2: BAZ2 suppresses G4 DNA formation. A-B.** Illustrative BG4 immunofluorescence of cells following a 2 hr treatment with DMSO, 5 μM PDS, 10 μM GSK2801 (**A**) or indicated siRNA for 48 hr (**B**). **C.** Quantitation of BRD9 by western blot following siRNA for (48 hr). **D-E.** Quantitation of cell cycle phases following 24 hr treatment with DMSO or 5 μM GSK2801 (**D**) or with indicated siRNA for 48 hr (**E**). ns, non-significant. *, $p < 0.05$; **, $p < 0.01$; ***, $p < 0.001$; ****, $p < 0.0001$. (Two-way ANOVA for multiple comparisons). Ct, Control; 2A, BAZ2A; 2B, BAZ2B.

### Supplementary Figure 3

A Replicates of MACS2 raw CUT&Tag

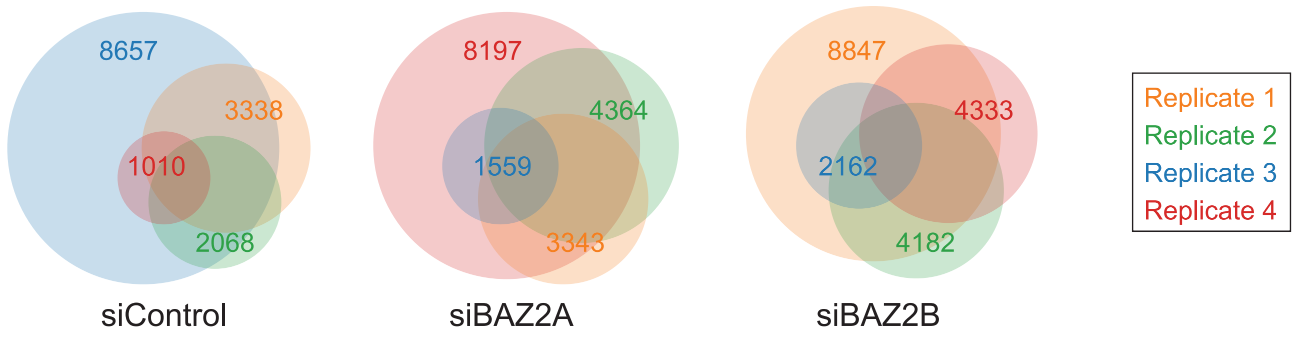


B SEACR with and without scaling

| | siCont | siBAZ2A | siBAZ2B |
|---|---|---|---|
| siCont | 10,605<br>10,605 | | |
| siBAZ2A | 8,625<br>8,625 | 11,861<br>11,861 | |
| siBAZ2B | 9,104<br>9,104 | 9,873<br>9,873 | 12,679<br>12,681 |

C

| Sample | % alignment to Drosophila |
|---|---|
| siControl-1 | 0.01 |
| siControl-2 | 0.01 |
| siControl-3 | 0.01 |
| siControl-4 | 0.01 |
| siBAZ2A-1 | 0.02 |
| siBAZ2A-2 | 0.01 |
| siBAZ2A-3 | 0.01 |
| siBAZ2A-4 | 0.01 |
| siBAZ2B-1 | 0.01 |
| siBAZ2B-2 | 0.02 |
| siBAZ2B-3 | 0.01 |
| siBAZ2B-4 | 0.02 |

**Supplementary Figure 3: Determination and normalization of high quality G4 CUT&Tag peaks. A.** Venn diagrams representing overlaps between the four replicates used to generate high-quality peaks after 48 hr of each indicated siRNA, total peaks for each replicate are noted. **B.** Number of high-quality peaks identified for each condition using SEACR with (green) and without (red) scaling to the spike-in reads. Diagonal is siCont, siBAZ2A, and siBAZ2B with other boxes showing overlap between conditions. **C.** Proportion of the total reads that aligned to the spike-in (*Drosophila*) genome for each replicate.

### Supplementary Figure 4

A

| siRNA | CUT&Tag | Mean peak length | G4s / 1000 bp |
|---|---|---|---|
| siCont | G4 DNA | 626 bp | 8.11 |
| siBAZ2A | | 577 | 8.28 |
| siBAZ2B | | 594 | 8.14 |
| siCont | H3K4me3 | 1,154 | 7.19 |
| Size-matched random control | | 626 | 1.10 |

B

| | Motif | E-value | Number of occurrences |
|---|---|---|---|
| siCont | 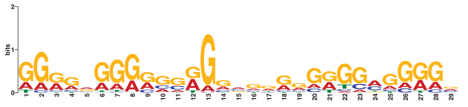 | 3.8e-038 | 1830 |
| | 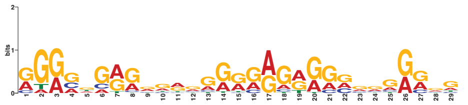 | 4.4e-041 | 1377 |
| siBAZ2A | 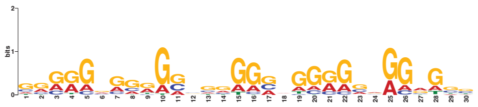 | 5.2e-068 | 3494 |
| | 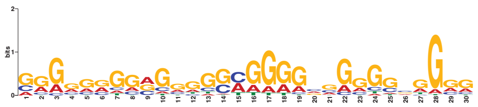 | 1.1e-034 | 2731 |
| siBAZ2B | 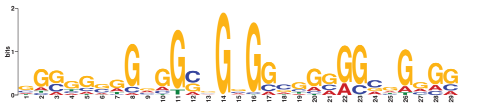 | 1.8e-041 | 2275 |
| | 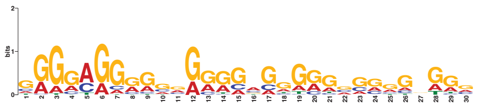 | 5.5e-043 | 2318 |

**Supplementary Figure 4: Characterization of G4 CUT&Tag peaks. A.** Mean peak length in base pair (bp) and number of G4 motifs per 1,000 bp for each condition as well as for a randomized control size-matched to the siControl G4 CUT&Tag peak dataset. **B.** Top two MEME Suite motifs identified in each condition of G4 CUT&Tag peaks.

### Supplementary Figure 5

A

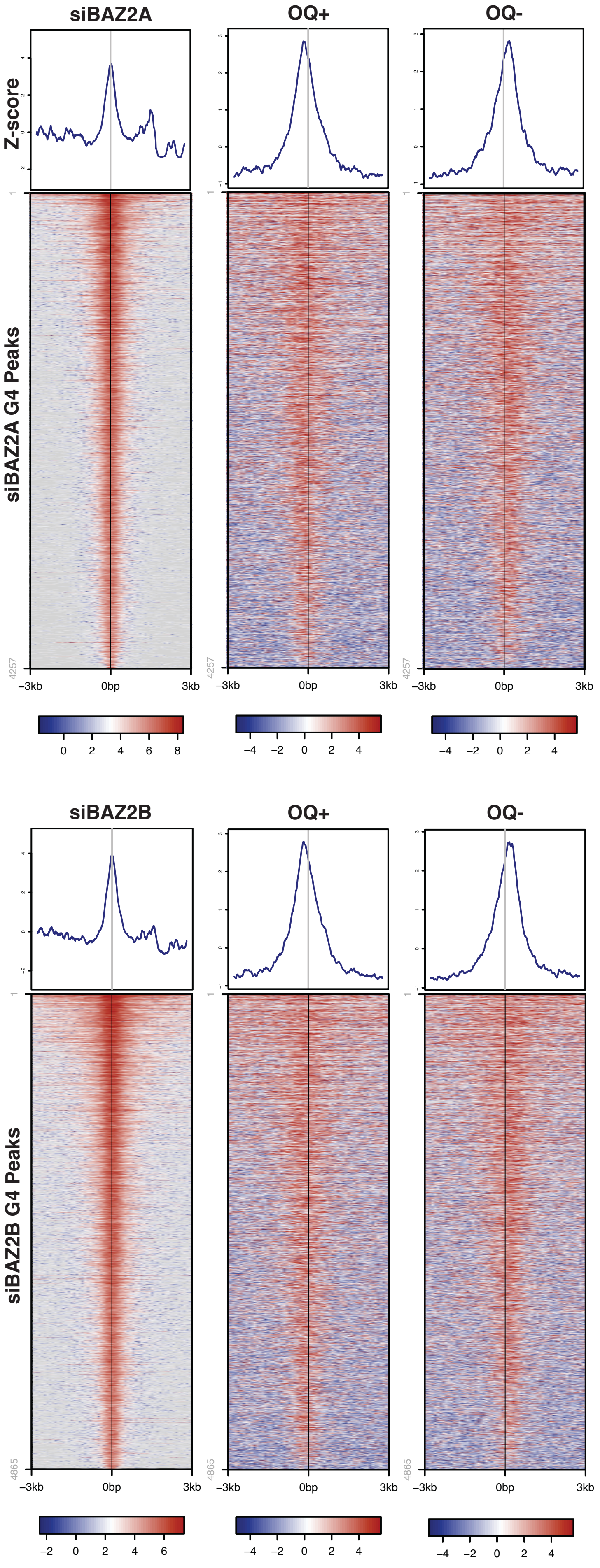


B

C

| | Peaks | Peaks with OQs | % peaks with OQs |
|---|---|---|---|
| siControl | 3332 | 2930 | 87.94 |
| siBAZ2A | 4257 | 3768 | 88.51 |
| siBAZ2B | 4865 | 4285 | 88.08 |

**Supplementary Figure 5: Concordance between G4 CUT&Tag peaks and Observed Quadruplexes. A-B.** G4 CUT&Tag and observed quadruplexes (OQ) with PDS (1) $\log_2$ transformed density heatmaps for peaks classified as G4 CUT&Tag in siBAZ2A (*n* = 4,257) (A) and siBAZ2B (*n* = 4,865)**. C.** Quantification of the overlap between G4 CUT&Tag peaks and OQs.

### Supplementary Figure 6

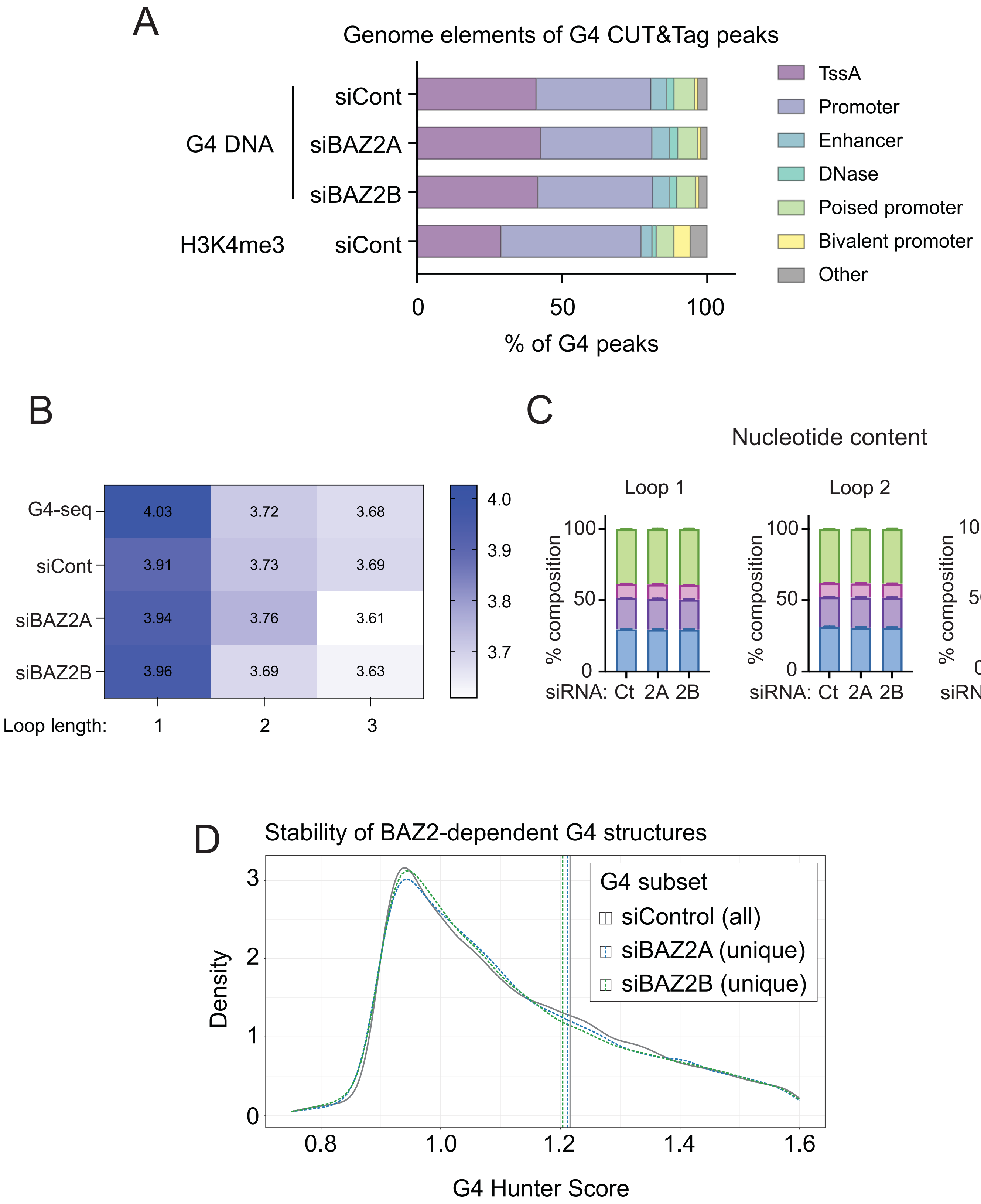

**Supplementary Figure 6: Characterization of G4 CUT&Tag peaks. A.** Distribution of G4 and H3K4me3 CUT&Tag peaks across regulatory elements derived from the Roadmap Epigenomics dataset in A549 cells. **B.** Mean loop length of the first three loops identified in G4 motifs (5' to 3') within G4 CUT&Tag peaks compared to G4-seq data (1). **C.** Nucleotide content in the first three loops. **D.** G4Hunter score distribution of G4 motifs found in all siControl peaks and the G4s unique to siBAZA or siBAZ2B (unique compared to siControl). The siBAZ2B distribution is significantly different from siControl ($p$ = 0.0016; Kolmogorov–Smirnov test). TssA, Transcription Start Site Active; siCont, siControl; Ct, Control; 2A, BAZ2A; 2B, BAZ2B

### Supplementary Figure 7

A

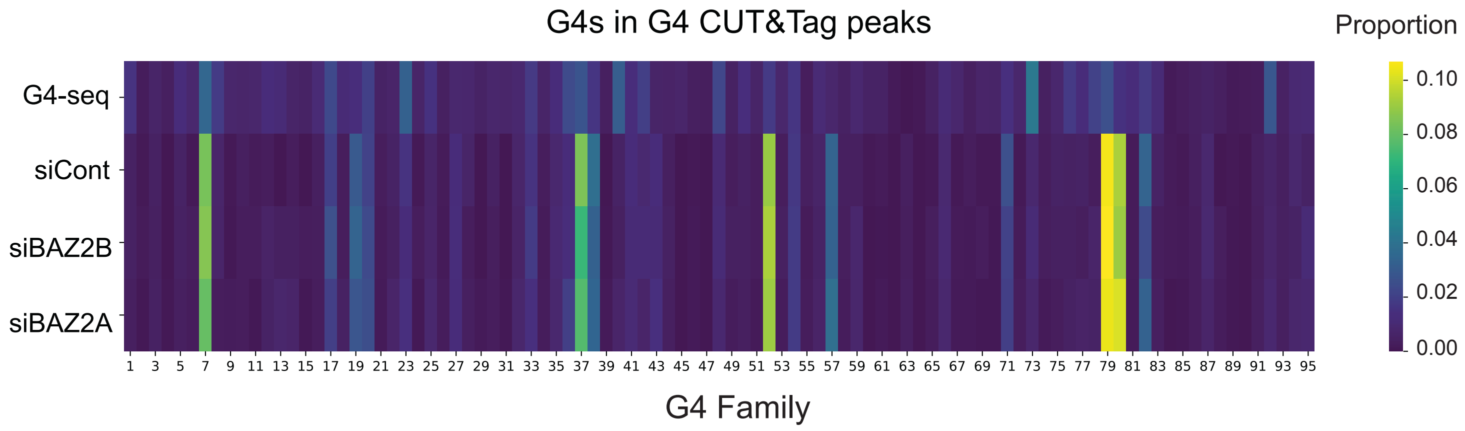


B

| Family | G repeat 1 | Loop 1 | G repeat 2 | Loop 2 | G repeat 3 | Loop 3 | G repeat 4 |
|---|---|---|---|---|---|---|---|
| 7 | GGG | C-CCSK | GGG | CDGS | GRGG | MR | GGG |
| 37 | GGG | SCA | GGG | CCA | GGG | CCA | GGG |
| 52 | GGG | SC- | GGG | C | GGG | C | GGG |
| 79 | GGG | M | GGG | -C | GGG | -- | GGG |
| 80 | GGG | -GC | GGG | SC---SS | GGG | GGM | GGG |

**Supplementary Figure 7: Clustering of G4 motifs. A.** G4 clustering analysis as performed by Neupane *et al.* (2) for G4 motifs identified by G4Catchall in the different G4 CUT&Tag peaks compared to those found in the G4-seq peaks. **B.** Attributes of the families of interest identified in the clustering analysis. siCont, siControl.

### Supplementary Figure 8

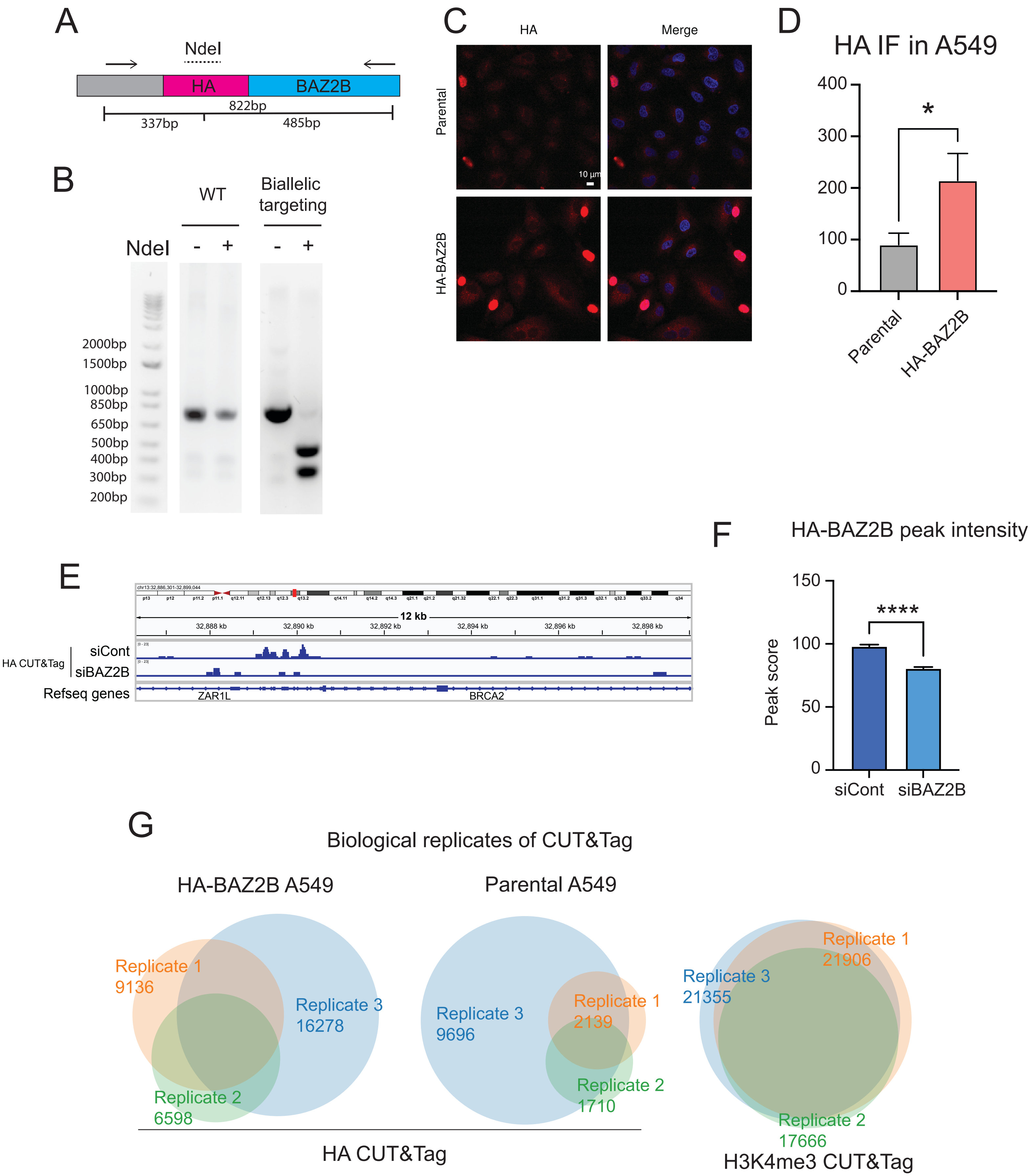

**Supplementary Figure 8: Insertion of an HA Tag at the Endogenous BAZ2B Locus. A-B.** Schematic (**A**) and PCR verification (**B**) of endogenous tagging of BAZ2B with an HA tag. Arrows indicate PCR primers. **C-D**. Illustrative immunofluorescence (**C**) and quantitation of HA IF signal (**D**). Blue in merged image is DAPI. **E.** Genome browser view at example locus showing HA-BAZ2B signal in A549 cells treated with either siControl (siCont) or siBAZ2B. **F**. Quantitation of HA-BAZ2B peak intensities at peaks common between siControl and siBAZ2B. **G**. Venn diagrams of overlap between replicates of HA CUT&Tag in the indicated cell lines. *, $p < 0.05$; ****, $p < 0.0001$ ($t$-test).

### Supplementary Figure 9

A

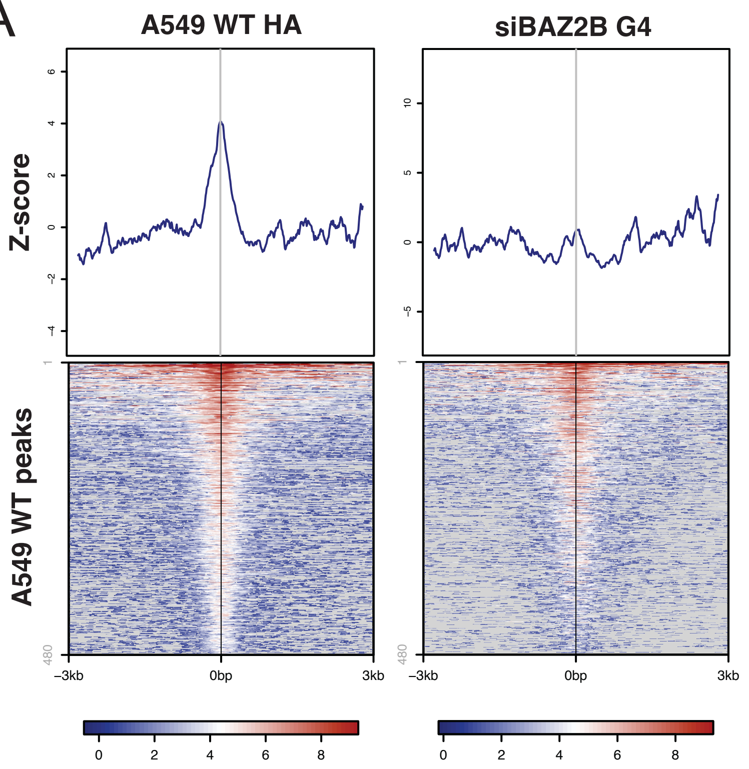


B

HA-BAZ2B CUT&Tag G4 enrichments

| Condition | Number of peaks | Mean peak length (bp) | % peaks containing G4s | Number of G4s / 1000 bp |
|---|---|---|---|---|
| HA-BAZ2B total peaks | 5043 | 1183.4 | 95.2 (44.3) | 7.4 (1.2) |
| HA-BAZ2B peaks overlapping with G4 CUT&Tag | 1947 | 1264.0 | 98.9 (46.6) | 7.8 (1.2) |
| HA-BAZ2B peaks not overlapping with G4 CUT&Tag | 3096 | 1132.8 | 93.0 (45.2) | 7.3 (1.3) |

C

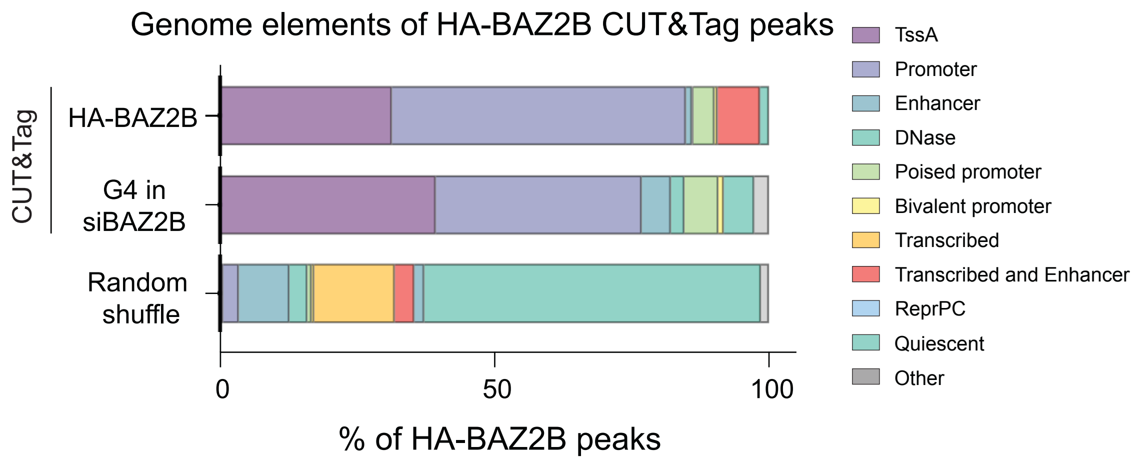


D

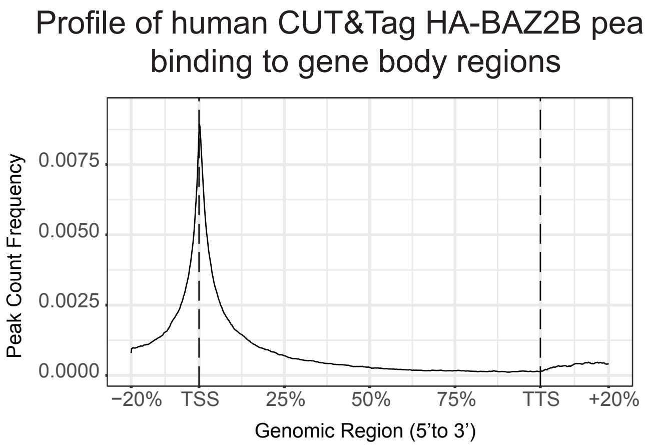


E

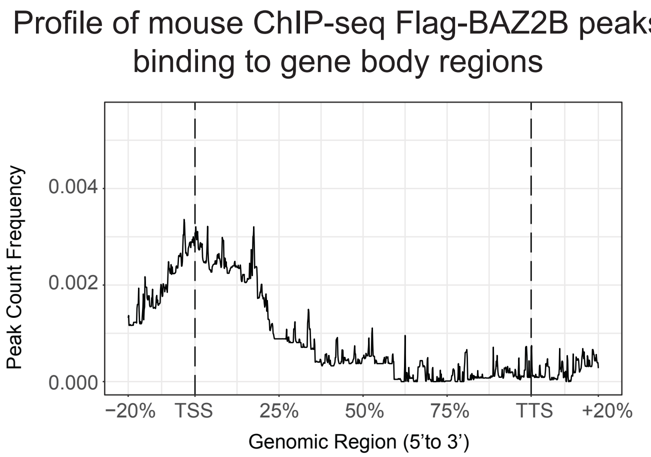

**Supplementary Figure 9: Characterization of HA-BAZ2B CUT&Tag peaks. A.** $Log_2$ transformed density heatmaps of HA CUT&Tag peaks in parental A549 cells (WT) and of siBAZ2B G4 CUT&Tag peaks within 10kb of parental HA CUT&Tag peaks (*n* = 480). **B**. G4 enrichment in HA-BAZ2B peaks. Red numbers in parentheses indicate the values found in size-matched random control datasets. **C**. Distribution of HA-BAZ2B peaks and a random size-matched control across regulatory elements derived from the Roadmap Epigenomics dataset in A549 cells. **D-E**. Average profile plots of HA-BAZ2B CUT&Tag peaks in gene body regions in A549 cells with endogenously tagged HA-BAZ2B (**D**) and Flag-Baz2b ChIP-seq peaks in gene body regions in H2.35 mouse liver cells with ectopic expression of Flag-Baz2b (**E**) from Jia *et al.* (3). WT, wild-type; TssA, Transcription Start Site Active; reprPC, repressed Polycomb states.

### Supplementary Figure 10

A

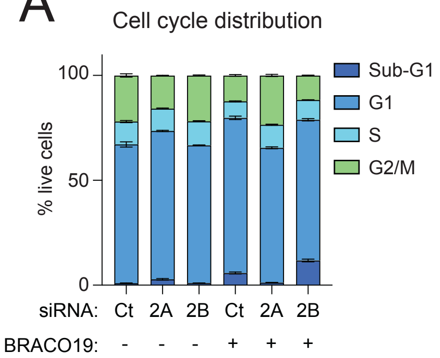


B

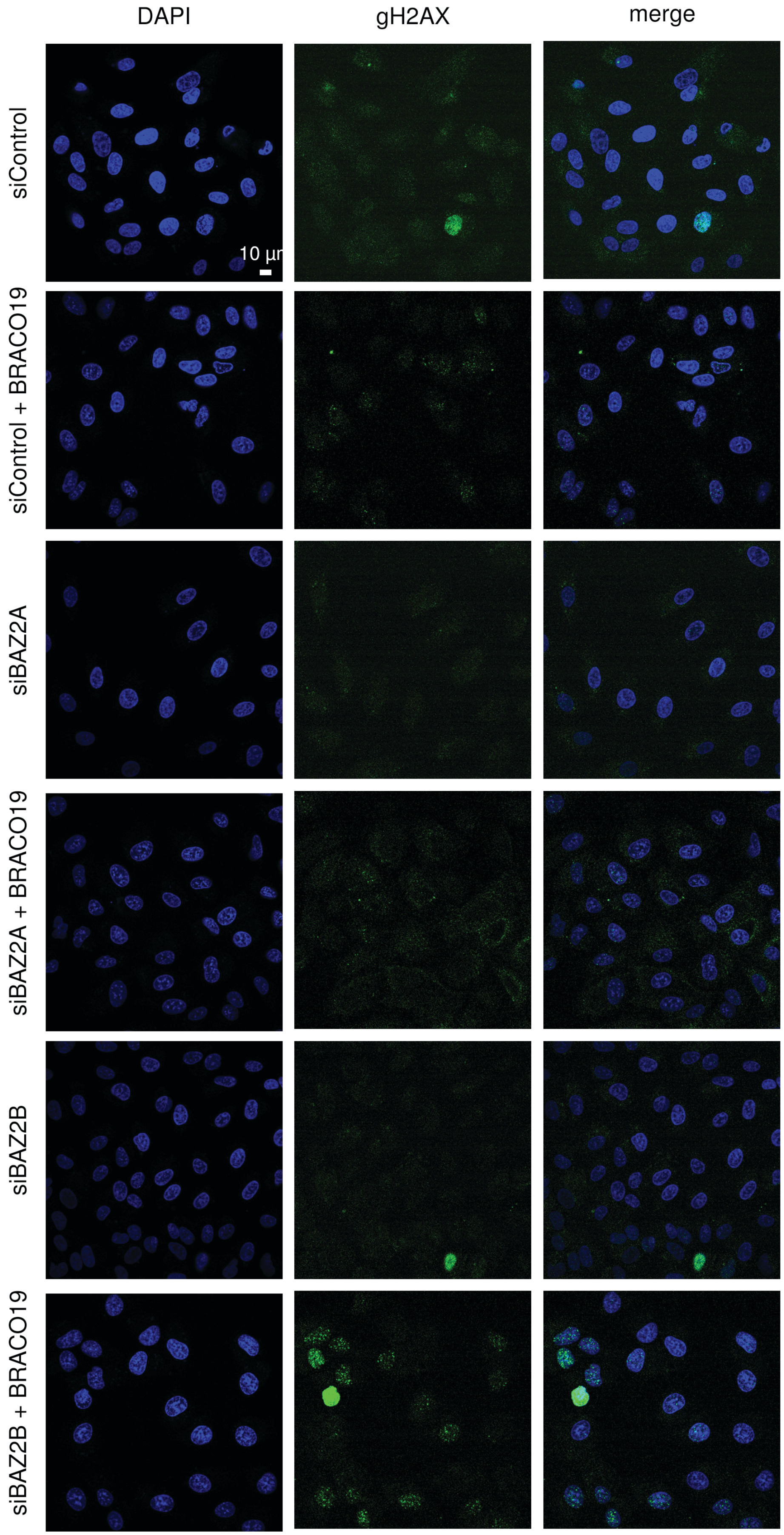

**Supplementary Figure 10: DNA damage in the absence of BAZ2. A**. Quantitation of cell cycle phase following 72 hr of the indicated siRNA treatments and 24 hr of 10 μM BRACO19. **B**. Illustrative γH2AX immunofluorescence (IF) of cells after 72 hr of the indicated siRNA and optional treatment with 10 μM BRACO19 for 24 hr.

### Supplementary Figure 11

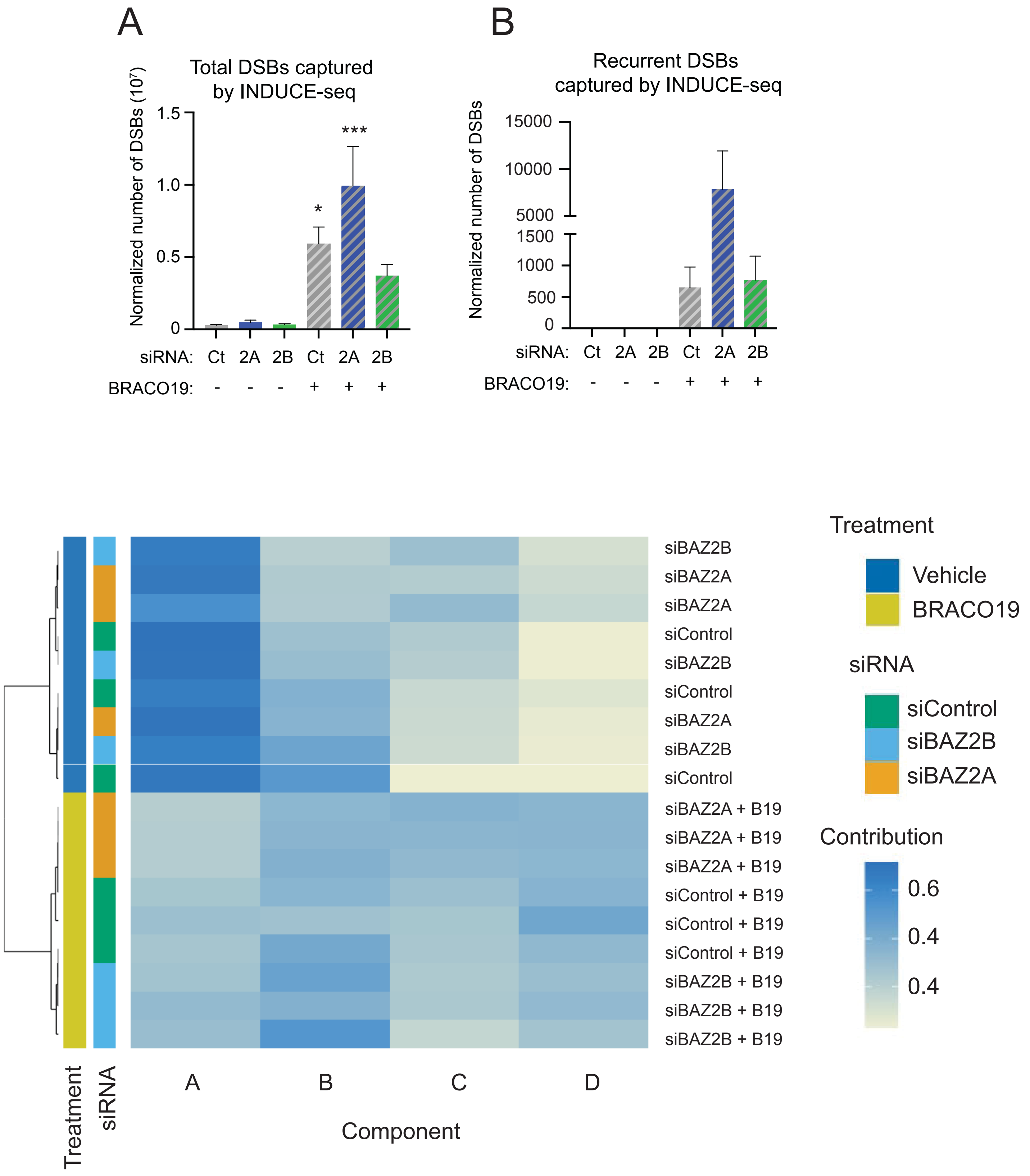

**Supplementary Figure 11: INDUCE-seq and NMF parameters. A-B**. Total (**A**) and recurrent (**B**) DSBs captured by INDUCE-seq normalized to the amount of extracted DNA ($n = 3$; *, $p < 0.05$; ****, $p < 0.0001$; one-way ANOVA for multiple comparisons). **C.** Heatmap showing hierarchal clustering of the contribution of NMF components to each INDUCE-seq sample, using cosine distance and Ward.D2 linkage method.

### Supplementary Figure 12

A

| GAT enrichments (1000 permutations) | | siCont | siBAZ2A | siBAZ2B | siCont +BRACO19 | siBAZ2A +BRACO19 | siBAZ2B +BRACO19 |
|---|---|---|---|---|---|---|---|
| All breaks | G4 peak overlap | 791 | 1536 | 1997 | 12,295 | 32,767 | 24,006 |
| | Expected | 644.7 | 1254.4 | 1559.2 | 13,099.6 | 34,856.7 | 21,837.3 |
| | Enrichment | 1.23 | 1.22 | 1.28 | 0.94 | 0.94 | 1.10 |
| | *p*-value | 0.001 | 0.001 | 0.001 | 0.001 | 0.001 | 0.001 |
| Recurrent breaks | G4 peak overlap | 0 | 0 | 0 | 193 | 1637 | 674 |
| | Expected | 0.008 | 0.002 | 0.017 | 48.46 | 586.54 | 125.70 |
| | Enrichment | 0.9921 | 0.998 | 0.9833 | 3.92 | 2.79 | 5.33 |
| | *p*-value | 0.996 | 0.999 | 0.994 | 0.001 | 0.001 | 0.001 |

**Supplementary Figure 12: DSB enrichment at G4 CUT&Tag peaks. A**. GAT analysis determining the enrichment of overlap between G4 CUT&Tag peaks and DSBs (*n* = 1,000 permutations).

Supplementary Figure 13

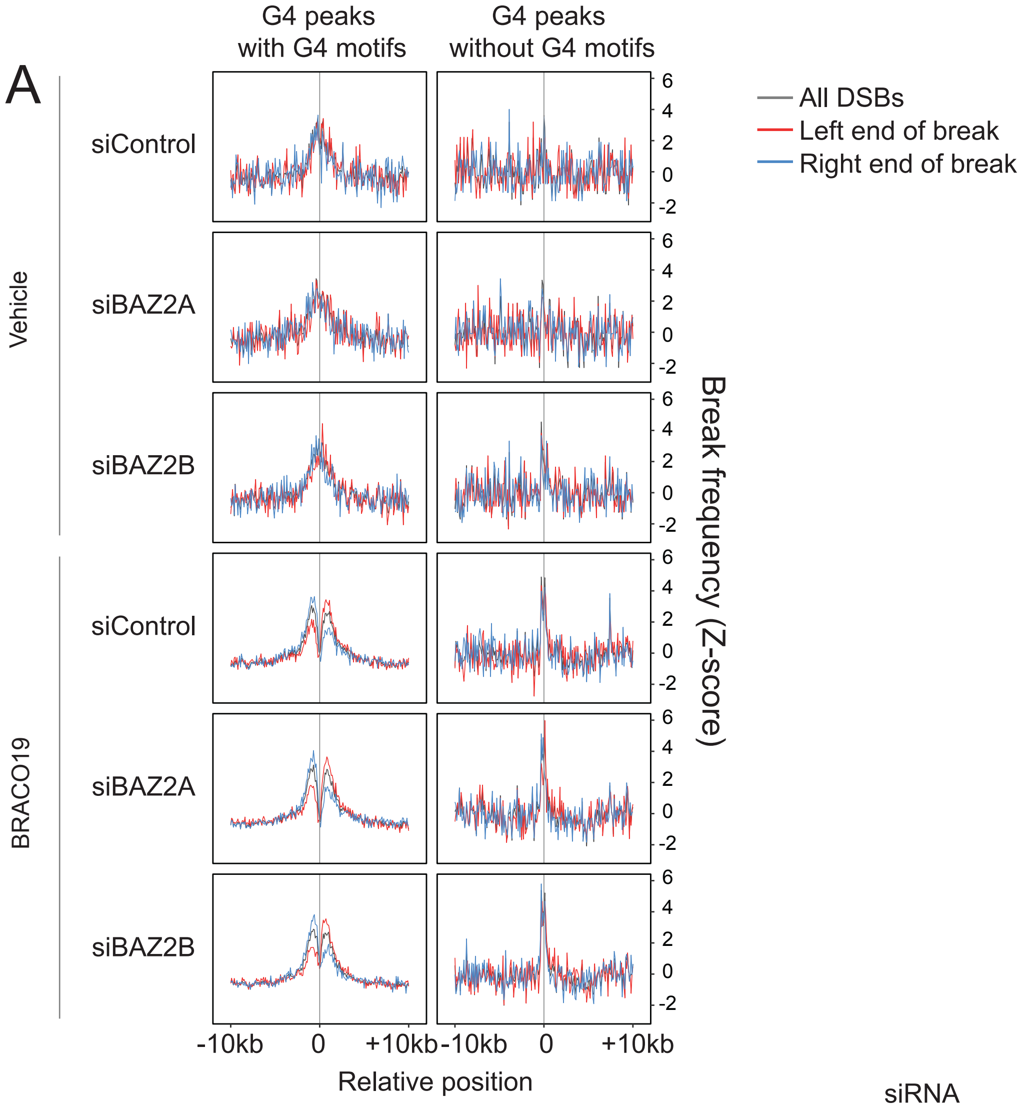


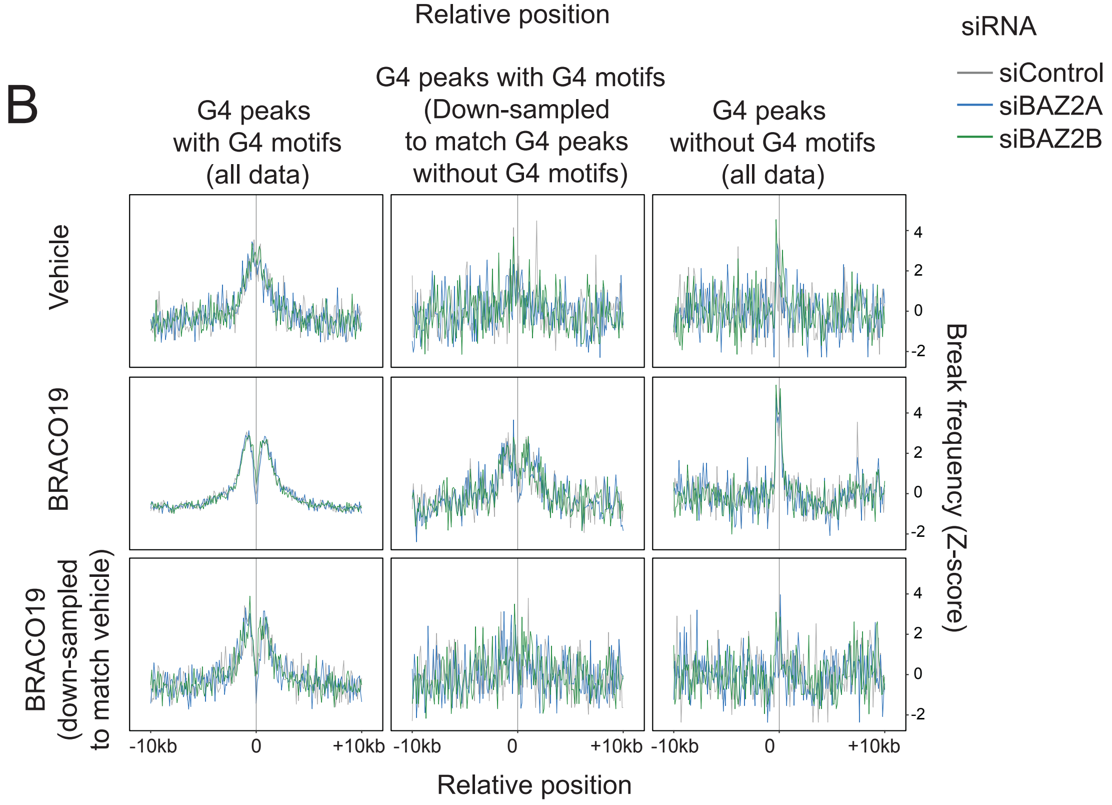

**Supplementary Figure 13: DSB distributions around G4 CUT&Tag peaks. A.** Average profile plots of DSB frequency (Z-score) within 10 kb of G4 CUT&Tag peaks with and without G4Catchall detected motifs with vehicle- or BRACO19-treatment; grey is all DSBs, red is the left end of DSBs and blue is the right end of DSBs. **B**. Profile plots of average DSB frequency (Z-score) within 10 kb of G4 CUT&Tag peaks with and without G4Catchall detected motifs with vehicle- or BRACO19-treatment. Data in middle column was down-sampled to match the number of G4 peaks without G4 motifs. Data in the bottom row were down-sampled to match the number of G4 peaks in the vehicle treatment; grey is siControl, blue is siBAZ2A and green is siBAZ2B.

### Supplementary Figure 14

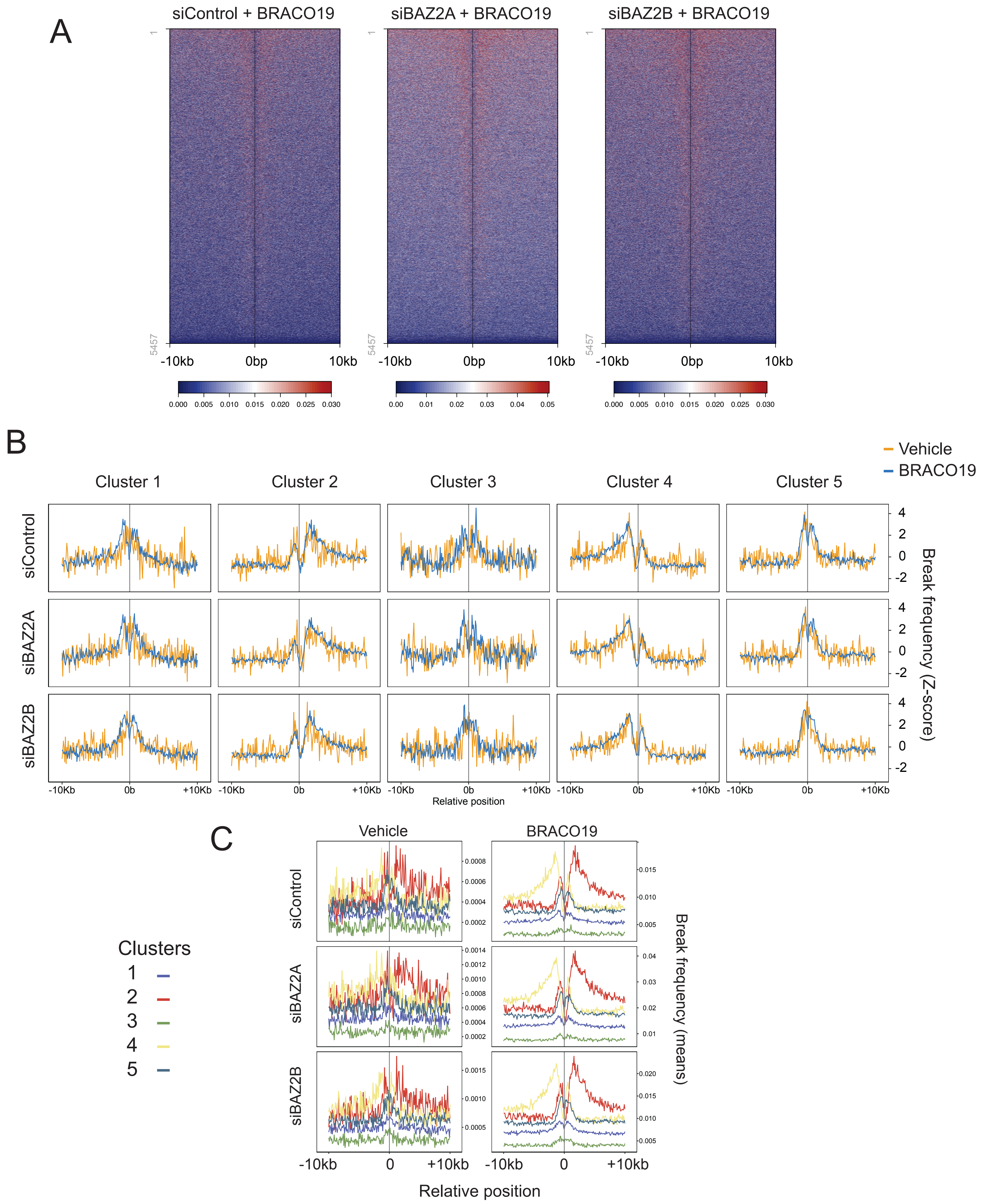

**Supplementary Figure 14: Clustering of DSB patterns surrounding G4 CUT&Tag peaks.**
**A**. Unclustered heatmaps of BRACO19-associated DSBs with indicated siRNA within 10 kb of G4 peaks (all conditions combined, *n* = 5,457). **B**. Profile plots of average DSB frequency (Z-score) within 10 kb of G4 CUT&Tag peaks sub-setted by cluster; yellow is vehicle-treated and blue is BRACO19-treated. **C**. Profile plots of average DSB frequency (non-normalized) to show the contribution of each cluster to the different conditions. Colors correspond to different clusters as indicated.

### Supplementary Figure 15

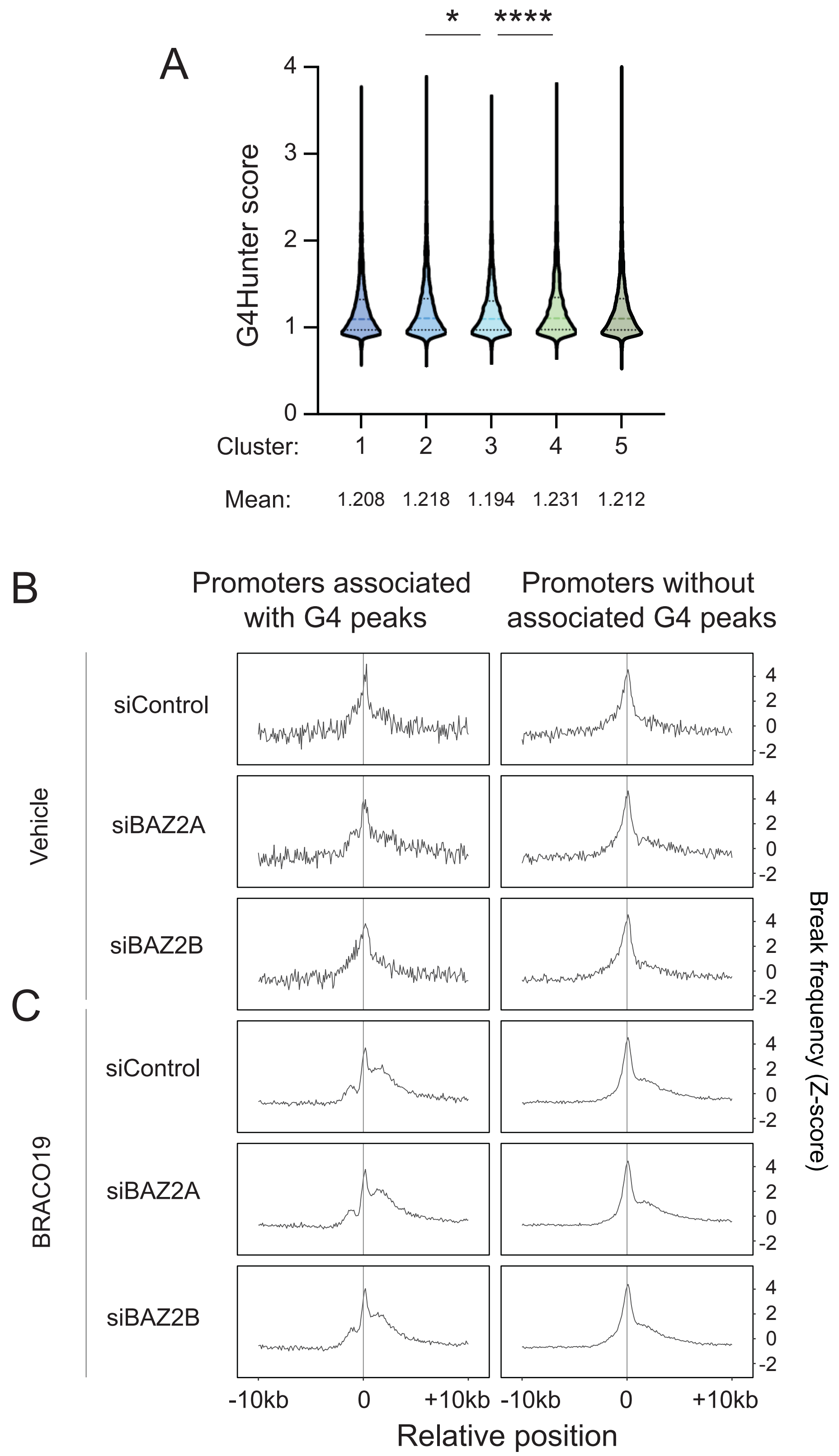

**Supplementary Figure 15: Characterization of clustering of DSB patterns. A**. G4Hunter score distribution for G4 motifs identified by G4Catchall in the peaks composing each cluster. **B-C**. Profile plots of average DSB frequency (Z-score) within 10 kb of promoters associated with G4 CUT&Tag peaks and promoters without associated G4 CUT&Tag peaks in vehicle- (**B**) or BRACO19-treated (**C**) samples. (*, $p < 0.05$; ****, $p < 0.0001$; one-way ANOVA for multiple comparisons).

### Supplementary Figure 16

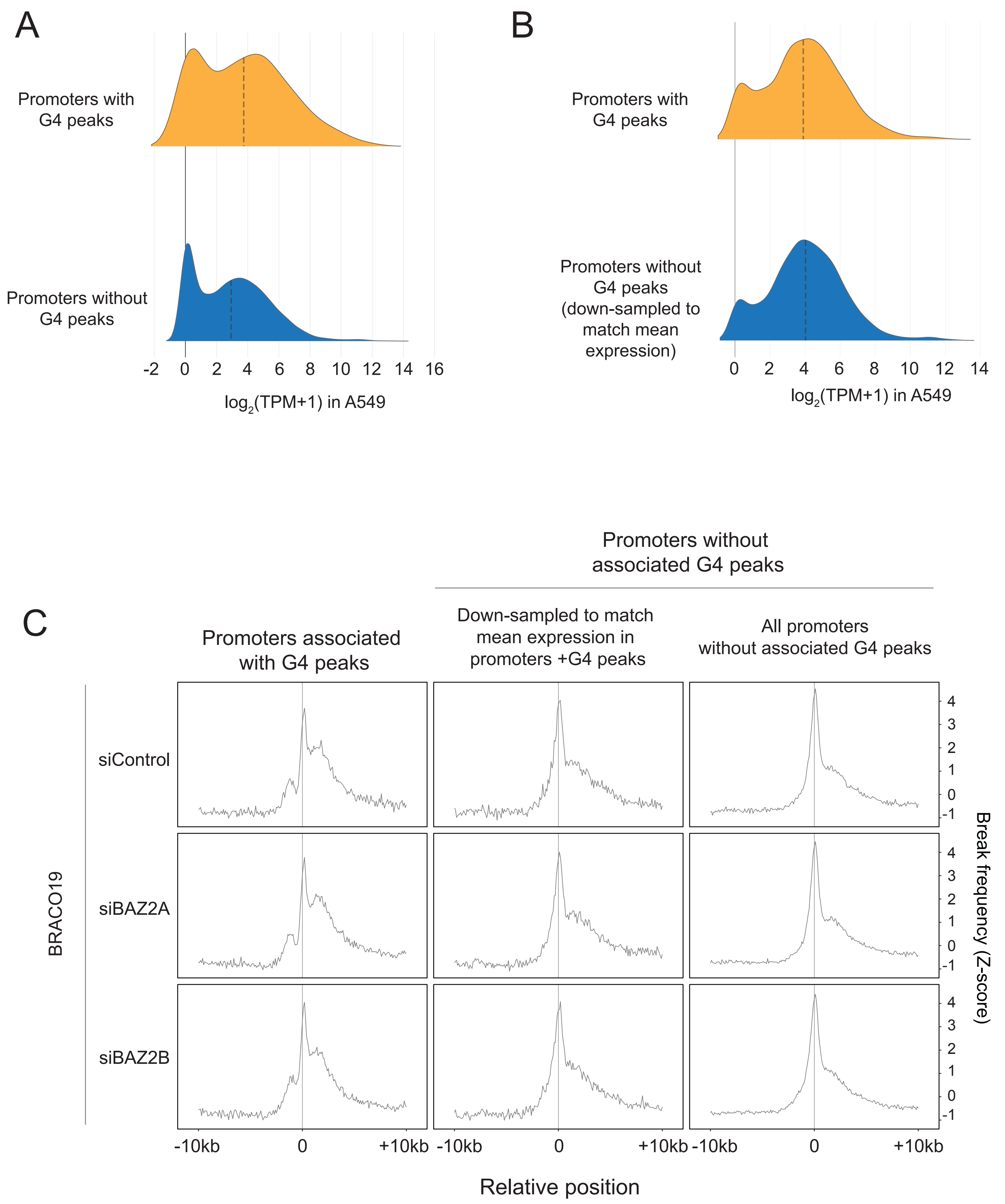

**Supplementary Figure 16: DSB distributions in the vicinity of promoters. A-B**. Gene expression levels from DepMap (4) of promoter-associated genes for promoters with and without G4 CUT&Tag peak overlap in the complete dataset (**A**) versus a promoter cohort without G4 CUT&Tag peak overlap, down-sampled to match the higher mean expression level in the G4 peak-overlapping set (**B**). **C.** Profile plots of average DSB frequency (Z-score) within 10 kb of promoters associated with G4 CUT&Tag peaks and promoters without associated G4 CUT&Tag. The middle column is downsampled to match the mean expression in promoters with overlapping G4 CUT&Tag peaks. TPM, transcripts per million. kb, kilobase.

### Supplementary Figure 17

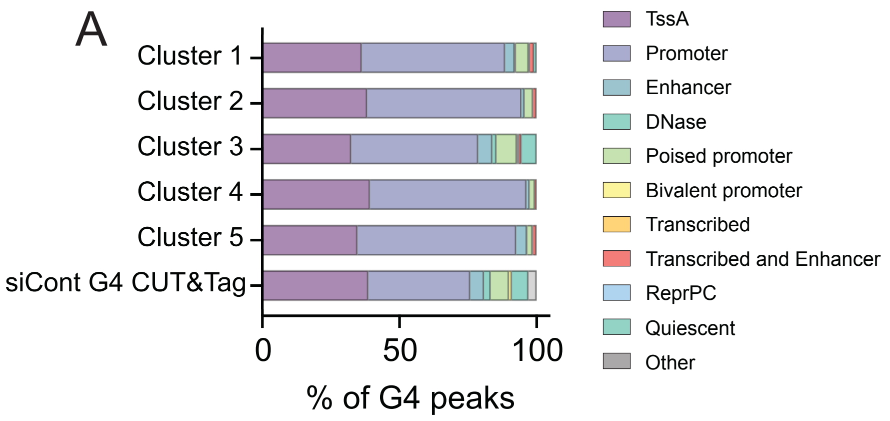


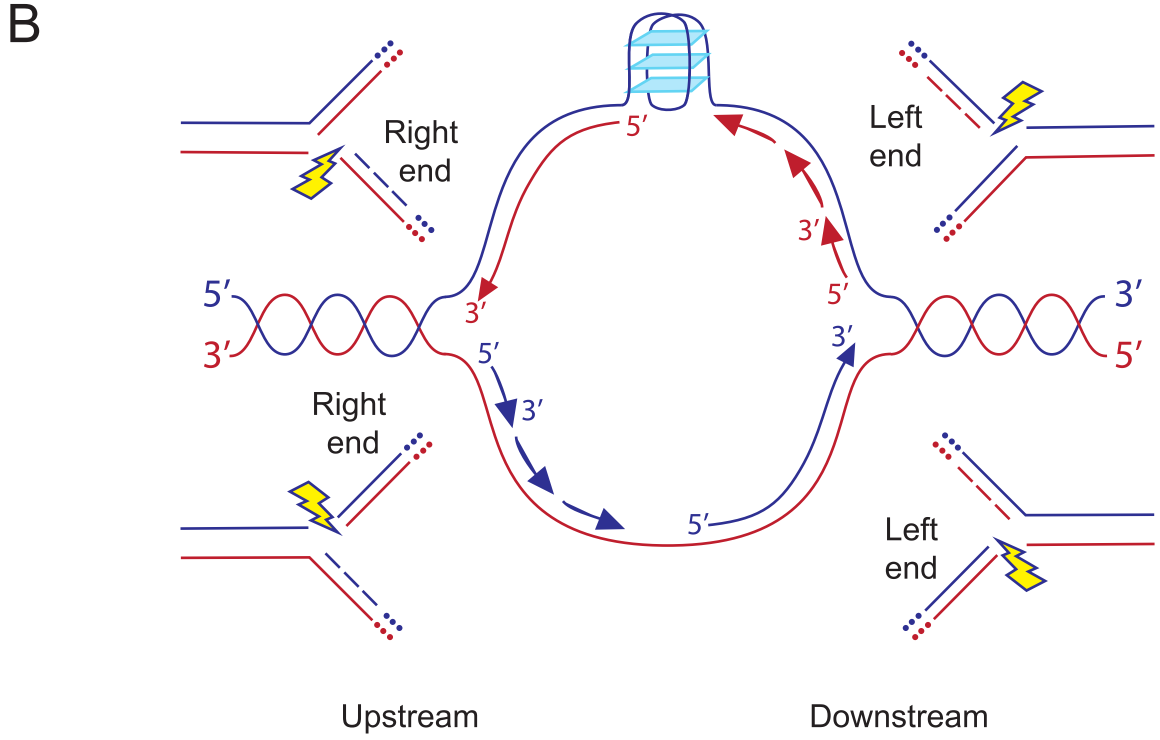

**Supplementary Figure 17: Genomic features may direct DSB in the vicinity of G4s. A.** Distribution of G4 peaks per cluster across regulatory elements derived from the Roadmap Epigenomics dataset in A549 cells compared to the siControl G4 CUT&Tag peak distribution. TssA, Transcription Start Site Active. **B**. Schematic representation of DSBs caused by fork collapse near origins of replication, which could hypothetically contain a G4 structure. Arrows indicate direction of replication. Model shows the preferential accumulation of right ends of one-sided DSBs upstream of the origin and accumulation of left ends of one-sided DSBs downstream of the origin.


1. G. Marsico *et al.*, Whole genome experimental maps of DNA G-quadruplexes in multiple species. *Nucleic Acids Res* **47**, 3862-3874 (2019).
2. A. Neupane, J. H. Chariker, E. C. Rouchka, Structural and Functional Classification of G-Quadruplex Families within the Human Genome. *Genes (Basel)* **14** (2023).
3. Y. Jia *et al.*, In vivo CRISPR screening identifies BAZ2 chromatin remodelers as druggable regulators of mammalian liver regeneration. *Cell Stem Cell* **29**, 372-385 e378 (2022).
4. A. Tsherniak *et al.*, Defining a Cancer Dependency Map. *Cell* **170**, 564-576 e516 (2017).

| Supplementary Table 2: Drugs used in experiments | | |
|---|---|---|
| **Name** | **Company** | **Catalog number** |
| (+)-JQ-1 | MedChem Express | HY-L005 |
| (+)-JQ1 PA | MedChem Express | HY-L005 |
| (R)-(-)-JQ1 Enantiomer | MedChem Express | HY-L005 |
| (S)-JQ-35 | MedChem Express | HY-L005 |
| 3-Aminobenzamide | MedChem Express | HY-L005 |
| 3-TYP | MedChem Express | HY-L005 |
| 5-Azacytidine | Millipore Sigma | A2385-100MG; HY-L005 |
| 6-Gingerol | MedChem Express | HY-L005 |
| 7-Methoxyisoflavone | MedChem Express | HY-L005 |
| A-366 | MedChem Express | HY-L005 |
| A-769662 | MedChem Express | HY-L005 |
| A-966492 | MedChem Express | HY-L005 |
| ABBV-744 | MedChem Express | HY-L005 |
| ACY-738 | MedChem Express | HY-L005 |
| AG14361 | MedChem Express | HY-L005 |
| AGK2 | MedChem Express | HY-L005 |
| AICAR | MedChem Express | HY-L005 |
| AICAR (phosphate) | MedChem Express | HY-L005 |
| AK-1 | MedChem Express | HY-L005 |
| AK-7 | MedChem Express | HY-L005 |
| AMG 900 | MedChem Express | HY-L005 |
| AMI-1 | MedChem Express | HY-L005 |
| AS8351 | MedChem Express | HY-L005 |
| AZD-1480 | MedChem Express | HY-L005 |
| AZD-2461 | MedChem Express | HY-L005 |
| AZD1152 | MedChem Express | HY-L005 |
| AZD5153 (6-Hydroxy-2-naphthoic acid) | MedChem Express | HY-100653A; HY-L005 |
| Alisertib | MedChem Express | HY-L005 |
| Amodiaquin (dihydrochloride dihydrate) | MedChem Express | HY-L005 |
| Anacardic Acid | MedChem Express | HY-L005 |
| Apabetalone | MedChem Express | HY-L005 |
| Atractylenolide I | MedChem Express | HY-L005 |
| BAY-299 | MedChem Express | HY-L005 |
| BAY1238097 | MedChem Express | HY-L005 |
| BCI-121 | MedChem Express | HY-L005 |
| BET bromodomain inhibitor | MedChem Express | HY-L005 |
| BG45 | MedChem Express | HY-L005 |
| BGP-15 | MedChem Express | HY-L005 |
| BI-7273 | MedChem Express | HY-L005 |
| BI-9564 | MedChem Express | HY-L005 |
| BML-210 | MedChem Express | HY-L005 |
| BMS-911543 | MedChem Express | HY-L005 |
| BRD4770 | MedChem Express | HY-L005 |

| | | |
|---|---|---|
| BRD73954 | MedChem Express | HY-L005 |
| BRD9539 | MedChem Express | HY-L005 |
| Barasertib-HQPA | MedChem Express | HY-L005 |
| Baricitinib | MedChem Express | HY-L005 |
| Baricitinib (phosphate) | MedChem Express | HY-L005 |
| Benzenebutyric acid | MedChem Express | HY-L005 |
| Birabresib | MedChem Express | HY-L005 |
| Bromosporine | MedChem Express | HY-L005 |
| Bufexamac | MedChem Express | HY-L005 |
| C-7280948 | MedChem Express | HY-15890; HY-L005 |
| CAY10602 | MedChem Express | HY-L005 |
| CCT129202 | MedChem Express | HY-L005 |
| CCT241736 | MedChem Express | HY-18161; HY-L005 |
| CHZ868 | MedChem Express | HY-L005 |
| CPI-169 racemate | MedChem Express | HY-L005 |
| CPI-203 | MedChem Express | HY-L005 |
| CPI-455 | MedChem Express | HY-L005 |
| CPI-637 | MedChem Express | HY-L005 |
| CUDC-101 | MedChem Express | HY-L005 |
| Cambinol | MedChem Express | HY-L005 |
| CeMMEC1 | MedChem Express | HY-L005 |
| Cerdulatinib | MedChem Express | HY-L005 |
| Citarinostat | MedChem Express | HY-L005 |
| Curcumin | MedChem Express | HY-L005 |
| DMSO | Fisher Scientific | BP231-100 |
| Daminozide | MedChem Express | HY-L005 |
| Danthron | MedChem Express | HY-L005 |
| Daphnetin | MedChem Express | HY-L005 |
| Decernotinib | MedChem Express | HY-L005 |
| Decitabine | MedChem Express | HY-L005 |
| Dihydrocoumarin | MedChem Express | HY-L005 |
| Domatinostat | MedChem Express | HY-L005 |
| Dorsomorphin (dihydrochloride) | MedChem Express | HY-L005 |
| Droxinostat | MedChem Express | HY-L005 |
| E7449 | MedChem Express | HY-L005 |
| EED inhibitor-1 | MedChem Express | HY-L005 |
| EED226 | MedChem Express | HY-L005 |
| EI1 | MedChem Express | HY-L005 |
| EPZ004777 | MedChem Express | HY-L005 |
| EPZ011989 | MedChem Express | HY-L005 |
| EPZ015866 | MedChem Express | HY-L005 |
| ETC-1002 | MedChem Express | HY-L005 |
| Entinostat | MedChem Express | HY-L005 |
| Enzastaurin | MedChem Express | HY-L005 |
| Filgotinib | MedChem Express | HY-L005 |

| | | |
|---|---|---|
| Fisetin | MedChem Express | HY-L005 |
| Flufenamic acid | MedChem Express | HY-L005 |
| GSK-5959 | MedChem Express | HY-L005 |
| GSK-J1 | MedChem Express | HY-L005 |
| GSK-J2 | MedChem Express | HY-L005 |
| GSK-J4 | MedChem Express | HY-L005 |
| GSK-LSD1 Dihydrochloride | MedChem Express | HY-L005 |
| GSK1324726A | MedChem Express | HY-L005 |
| GSK2801 | MedChem Express, Sigma-Aldrich | HY-15658; HY-L005; SML0768-5MG |
| GSK2879552 | MedChem Express | HY-18632 |
| GSK3326595 | MedChem Express | HY-L005 |
| GSK484 | MedChem Express | HY-L005 |
| GSK503 | MedChem Express | HY-L005 |
| GSK6853 | MedChem Express | HY-L005 |
| Ginkgolide C | MedChem Express | HY-L005 |
| Go 6983 | MedChem Express | HY-L005 |
| HA-100 | MedChem Express | HY-L005 |
| HDAC8-IN-1 | MedChem Express | HY-L005 |
| HLCL-61 (hydrochloride) | MedChem Express | HY-L005 |
| HPOB | MedChem Express | HY-L005 |
| Hinokitiol | Millipore Sigma | 469521-5G; HY-L005 |
| Histone Acetyltransferase Inhibitor II | MedChem Express | HY-L005 |
| I-BET151 | MedChem Express | HY-L005 |
| I-BRD9 | MedChem Express | HY-L005 |
| IOX1 | MedChem Express | HY-L005 |
| ITSA-1 | MedChem Express | HY-L005 |
| Inauhzin | MedChem Express | HY-L005 |
| Ingenol | MedChem Express | HY-L005 |
| Iniparib | MedChem Express | HY-L005 |
| JANEX-1 | MedChem Express | HY-L005 |
| JIB-04 | MedChem Express | HY-L005 |
| JNJ-7706621 | MedChem Express | HY-L005 |
| JQ-1 (carboxylic acid) | MedChem Express | HY-L005 |
| JW 55 | MedChem Express | HY-L005 |
| KG-501 | MedChem Express | HY-L005 |
| KW-2449 | MedChem Express | HY-L005 |
| LFM-A13 | MedChem Express | HY-L005 |
| LMK-235 | MedChem Express | HY-L005 |
| Lomeguatrib | MedChem Express | HY-L005 |
| MC1568 | MedChem Express | HY-L005 |
| ME0328 | MedChem Express | HY-L005 |
| MG 149 | MedChem Express | HY-L005 |
| MI-2 | MedChem Express | HY-15222; HY-L005 |
| MI-3 | MedChem Express | HY-L005 |

| | | |
|---|---|---|
| MK-8745 | MedChem Express | HY-L005 |
| ML324 | MedChem Express | HY-L005 |
| MLN8054 | MedChem Express | HY-L005 |
| MM-102 (TFA) | MedChem Express | HY-L005 |
| MN-64 | MedChem Express | HY-L005 |
| MS023 | MedChem Express | HY-L005 |
| MS049 | MedChem Express | HY-L005 |
| MS436 | MedChem Express | HY-L005 |
| Metformin (hydrochloride) | MedChem Express | HY-L005 |
| Mitoxantrone | MedChem Express | HY-L005 |
| Mitoxantrone (dihydrochloride) | MedChem Express | HY-L005 |
| Mivebresib | MedChem Express | HY-L005 |
| Molibresib | MedChem Express | HY-L005 |
| NI-57 | MedChem Express | HY-L005 |
| NKL 22 | MedChem Express | HY-L005 |
| NMS-P118 | MedChem Express | HY-L005 |
| Nicotinamide | MedChem Express | HY-L005 |
| Niraparib | MedChem Express | HY-L005 |
| Niraparib tosylate | MedChem Express | HY-L005 |
| O-304 | MedChem Express | HY-112233; HY-L005 |
| OF-1 | MedChem Express | HY-L005 |
| OICR-9429 | MedChem Express | HY-L005 |
| ORY-1001(trans) | MedChem Express | HY-L005 |
| OSS_128167 | MedChem Express | HY-L005 |
| Oclacitinib (maleate) | MedChem Express | HY-L005 |
| Olaparib | MedChem Express | HY-L005 |
| PCI-34051 | MedChem Express | HY-L005 |
| PF-06409577 | MedChem Express | HY-L005 |
| PF-06651600 | MedChem Express | HY-L005 |
| PF-06726304 | MedChem Express | HY-L005 |
| PF-CBP1 hydrochloride | MedChem Express | HY-L005 |
| PFI-1 | MedChem Express | HY-L005 |
| PFI-2 (hydrochloride) | MedChem Express | HY-L005 |
| PFI-3 | MedChem Express | HY-L005 |
| PFI-4 | MedChem Express | HY-L005 |
| PHA-680632 | MedChem Express | HY-L005 |
| PJ34 | MedChem Express | HY-L005 |
| PJ34 (hydrochloride) | MedChem Express | HY-L005 |
| PLX51107 | MedChem Express | HY-L005 |
| PTACH | MedChem Express | HY-L005 |
| Pacritinib | MedChem Express | HY-L005 |
| Pamiparib | MedChem Express | HY-L005 |
| Peficitinib | MedChem Express | HY-L005 |
| Phenformin (hydrochloride) | MedChem Express | HY-L005 |
| Pimelic Diphenylamide 106 | MedChem Express | HY-L005 |

| | | |
|---|---|---|
| Pinometostat | MedChem Express | HY-L005 |
| Pyridostatin | Millipore Sigma | SML2690-25MG |
| RG108 | MedChem Express | HY-L005 |
| RG2833 | MedChem Express | HY-L005 |
| RGFP966 | MedChem Express | HY-L005 |
| Remetinostat | MedChem Express | HY-L005 |
| Remodelin (hydrobromide) | MedChem Express | HY-L005 |
| Resminostat (hydrochloride) | MedChem Express | HY-L005 |
| Ricolinostat | MedChem Express | HY-L005 |
| Ro 31-8220 (mesylate) | MedChem Express | HY-L005 |
| Rucaparib (phosphate) | MedChem Express | HY-L005 |
| Ruxolitinib | MedChem Express | HY-L005 |
| Ruxolitinib (phosphate) | MedChem Express | HY-L005 |
| SAR-20347 | MedChem Express | HY-L005 |
| SGC-CBP30 | MedChem Express | HY-L005 |
| SGC0946 | MedChem Express | HY-L005 |
| SGC707 | MedChem Express | HY-L005 |
| SID 3712249 | MedChem Express | HY-L005 |
| SNS-314 | MedChem Express | HY-L005 |
| SP2509 | MedChem Express | HY-L005 |
| SR-4370 | MedChem Express | HY-L005 |
| Salermide | MedChem Express | HY-L005 |
| Santacruzamate A | MedChem Express | HY-L005 |
| Scriptaid | MedChem Express | HY-L005 |
| Sinapinic acid | MedChem Express | HY-L005 |
| Sirtinol | MedChem Express | HY-L005 |
| Sodium Butyrate | MedChem Express | HY-L005 |
| Sodium phenylbutyrate | MedChem Express | HY-L005 |
| Solcitinib | MedChem Express | HY-L005 |
| Sotrastaurin | MedChem Express | HY-L005 |
| Sulforaphane | MedChem Express | HY-L005 |
| TAK-632 | MedChem Express | HY-L005 |
| TAS-301 | MedChem Express | HY-L005 |
| TMP195 | MedChem Express | HY-L005 |
| TMP269 | MedChem Express | HY-L005 |
| TMPyP4 | Sigma-Aldrich | 613560-25MG |
| Tacedinaline | MedChem Express | HY-50934; HY-L005 |
| Talazoparib tosylate | MedChem Express | HY-108413; HY-L005 |
| Tasquinimod | MedChem Express | HY-L005 |
| Tazemetostat | MedChem Express | HY-L005 |
| Tenovin-1 | MedChem Express | HY-L005 |
| Tofacitinib | MedChem Express | HY-L005 |
| Tofacitinib (citrate) | MedChem Express | HY-L005 |
| Tozasertib | MedChem Express | HY-L005 |
| Tranylcypromine (hemisulfate) | MedChem Express | HY-L005 |

| | | |
|---|---|---|
| Tubastatin A (Hydrochloride) | MedChem Express | HY-L005 |
| Tubastatin-A | MedChem Express | HY-L005 |
| Tucidinostat | MedChem Express | HY-L005 |
| UF010 | MedChem Express | HY-L005 |
| UNC 669 | MedChem Express | HY-L005 |
| UNC0224 | MedChem Express | HY-L005 |
| UNC0321 | MedChem Express | HY-L005 |
| UNC0379 | MedChem Express | HY-L005 |
| UNC1215 | MedChem Express | HY-L005 |
| UNC3866 | MedChem Express | HY-L005 |
| UPF 1069 | MedChem Express | HY-L005 |
| Valproic acid | MedChem Express | HY-L005 |
| Valproic acid (sodium salt) | MedChem Express | HY-L005 |
| Valrubicin | MedChem Express | HY-L005 |
| Veliparib | MedChem Express | HY-10129; HY-L005 |
| Veliparib (dihydrochloride) | MedChem Express | HY-L005 |
| Verbascoside | MedChem Express | HY-L005 |
| Vorinostat | MedChem Express | HY-L005 |
| WDR5-0103 | MedChem Express | HY-L005 |
| WHI-P154 | MedChem Express | HY-L005 |
| WHI-P97 | MedChem Express | HY-L005 |
| WM-1119 | MedChem Express | HY-L005 |
| WT-161 | MedChem Express | HY-L005 |
| WZ4003 | MedChem Express | HY-L005 |
| XAV-939 | MedChem Express | HY-L005 |
| XL228 | MedChem Express | HY-L005 |
| XY1 | MedChem Express | HY-L005 |
| ZL0420 | MedChem Express | HY-112149; HY-L005 |
| ZLN024 (hydrochloride) | MedChem Express | HY-L005 |
| ZM-447439 | MedChem Express | HY-L005 |
| ZM39923 (hydrochloride) | MedChem Express | HY-L005 |
| Zebularine | MedChem Express | HY-L005 |
| dBET1 | MedChem Express | HY-L005 |
| y-Oryzanol | MedChem Express | HY-L005 |
| BRACO19 | Millipore Sigma | SML0560-5MG |
| CX5461 | Sigma-Aldrich | 5092650001 |

| Supplementary Table 2: Oligonucleotide sequences | | | |
|---|---|---|---|
| **siRNA** | **Sequences** | **Company** | **Catalog number** |
| Human siControl (Non-targetimg pool) | UGGUUUACAUGUCGACUAA, UGGUUUACAUGUUGUGUGA, UGGUUUACAUGUUUUCUGA, UGGUUUACAUGUUUUCCUA | Dharmacon | D-001810-10-20 |
| Human siBAZ2A (smartpool) | GAUAAGACACCAAGACGUA, CGUCGAGGAUUCAGGCAAU, GUGGAGGACUAUAUGGUAU, GGGAGACGGCGUUGGGAUA | Dharmacon | L-020470-01-0010 |
| Human siBAZ2B (smartpool) | AGACAAUGUUUCCGAGAUU, CAGGAUGAGACGUCGGAAA, CCAAGUAACUCGAGAUAUU, CGACAAGAACAUUGAUUAU | Dharmacon | L-020487-01-0010 |
| Human siBRD9 (siGENOME) | CAAUGAAGAUACAGCUGUU | Dharmacon | D-014250-01-0002 |
| **Primers** | **Sequences** | **Company** | **Experiment** |
| hBactin FOR | CACCAACTGGGACGACAT | Thermo Fisher | qRT-PCR |
| hBactin REV | ACAGCCTGGATAGCAACG | Thermo Fisher | qRT-PCR |
| hBAZ2A FOR | AAGATGTGTGGCTACAATGG | Thermo Fisher | qRT-PCR |
| hBAZ2A REV | TCTGCACCCATCAGCTCCG | Thermo Fisher | qRT-PCR |
| hBAZ2B FOR | GTGGCTTCAGTAGTTTCAAAAGG | Thermo Fisher | qRT-PCR |
| hBAZ2B REV | GACACTGTGGACAGGTTAAACG | Thermo Fisher | qRT-PCR |
| hBRD9 FOR | CCAGCTGAAGCAGAGAAGAAA | Thermo Fisher | qRT-PCR |
| hBRD9 REV | CGACATGAACTCCAGAACAGAG | Thermo Fisher | qRT-PCR |
| BAZ2B 502 FOR | CAGGCCAGATTCTTGGACACT | Thermo Fisher | PCR |
| BAZ2B Flag REV | CCAGTGCTATCCCAGCGTAA | Thermo Fisher | PCR |
| **i5 adaptors** | **Sequences** | **Company** | **Experiment** |
| v2_Ad1.1_TAGATCGC | AATGATACGGCGACCACCGAGATCTACACTAGATCGCTCGTCGGCAGCGTCAGATGTGTAT | Thermo Fisher | CUT&Tag library prep |
| v2_Ad1.2_CTCTCTAT | AATGATACGGCGACCACCGAGATCTACACCTCTCTATTCGTCGGCAGCGTCAGATGTGTAT | Thermo Fisher | CUT&Tag library prep |
| v2_Ad1.3_TATCCTCT | AATGATACGGCGACCACCGAGATCTACACTATCCTCTTCGTCGGCAGCGTCAGATGTGTAT | Thermo Fisher | CUT&Tag library prep |
| v2_Ad1.4_AGAGTAGA | AATGATACGGCGACCACCGAGATCTACACAGAGTAGATCGTCGGCAGCGTCAGATGTGTAT | Thermo Fisher | CUT&Tag library prep |
| v2_Ad1.5_GTAAGGAG | AATGATACGGCGACCACCGAGATCTACACGTAAGGAGTCGTCGGCAGCGTCAGATGTGTAT | Thermo Fisher | CUT&Tag library prep |
| v2_Ad1.6_ACTGCATA | AATGATACGGCGACCACCGAGATCTACACACTGCATATCGTCGGCAGCGTCAGATGTGTAT | Thermo Fisher | CUT&Tag library prep |
| v2_Ad1.7_AAGGAGTA | AATGATACGGCGACCACCGAGATCTACACAAGGAGTATCGTCGGCAGCGTCAGATGTGTAT | Thermo Fisher | CUT&Tag library prep |
| v2_Ad1.8_CTAAGCCT | AATGATACGGCGACCACCGAGATCTACACCTAAGCCTTCGTCGGCAGCGTCAGATGTGTAT | Thermo Fisher | CUT&Tag library prep |
| v2_Ad1.9_TGGAAATC | AATGATACGGCGACCACCGAGATCTACACTGGAAATCTCGTCGGCAGCGTCAGATGTGTAT | Thermo Fisher | CUT&Tag library prep |
| v2_Ad1.10_AACATGAT | AATGATACGGCGACCACCGAGATCTACACAACATGATTCGTCGGCAGCGTCAGATGTGTAT | Thermo Fisher | CUT&Tag library prep |
| v2_Ad1.11_TGATGAAA | AATGATACGGCGACCACCGAGATCTACACTGATGAAATCGTCGGCAGCGTCAGATGTGTAT | Thermo Fisher | CUT&Tag library prep |
| v2_Ad1.12_GTCGGACT | AATGATACGGCGACCACCGAGATCTACACGTCGGACTTCGTCGGCAGCGTCAGATGTGTAT | Thermo Fisher | CUT&Tag library prep |
| v2_Ad1.13_TTTCTAGC | AATGATACGGCGACCACCGAGATCTACACTTTCTAGCTCGTCGGCAGCGTCAGATGTGTAT | Thermo Fisher | CUT&Tag library prep |
| v2_Ad1.14_TAACCAAG | AATGATACGGCGACCACCGAGATCTACACTAACCAAGTCGTCGGCAGCGTCAGATGTGTAT | Thermo Fisher | CUT&Tag library prep |
| v2_Ad1.15_GTGTATCG | AATGATACGGCGACCACCGAGATCTACACGTGTATCGTCGTCGGCAGCGTCAGATGTGTAT | Thermo Fisher | CUT&Tag library prep |
| v2_Ad1.16_TCCATCAA | AATGATACGGCGACCACCGAGATCTACACTCCATCAATCGTCGGCAGCGTCAGATGTGTAT | Thermo Fisher | CUT&Tag library prep |
| **i7 adaptors** | **Sequences** | **Company** | **Experiment** |
| v2_Ad2.1_TAAGGCGA | CAAGCAGAAGACGGCATACGAGATTCGCCTTAGTCTCGTGGGCTCGGAGATGTG | Thermo Fisher | CUT&Tag library prep |
| v2_Ad2.2_CGTACTAG | CAAGCAGAAGACGGCATACGAGATCTAGTACGGTCTCGTGGGCTCGGAGATGTG | Thermo Fisher | CUT&Tag library prep |
| v2_Ad2.3_AGGCAGAA | CAAGCAGAAGACGGCATACGAGATTTCTGCCTGTCTCGTGGGCTCGGAGATGTG | Thermo Fisher | CUT&Tag library prep |
| v2_Ad2.4_TCCTGAGC | CAAGCAGAAGACGGCATACGAGATGCTCAGGAGTCTCGTGGGCTCGGAGATGTG | Thermo Fisher | CUT&Tag library prep |
| v2_Ad2.5_GGACTCCT | CAAGCAGAAGACGGCATACGAGATAGGAGTCCGTCTCGTGGGCTCGGAGATGTG | Thermo Fisher | CUT&Tag library prep |
| v2_Ad2.6_TAGGCATG | CAAGCAGAAGACGGCATACGAGATCATGCCTAGTCTCGTGGGCTCGGAGATGTG | Thermo Fisher | CUT&Tag library prep |
| v2_Ad2.7_CTCTCTAC | CAAGCAGAAGACGGCATACGAGATGTAGAGAGGTCTCGTGGGCTCGGAGATGTG | Thermo Fisher | CUT&Tag library prep |
| v2_Ad2.8_CAGAGAGG | CAAGCAGAAGACGGCATACGAGATCCTCTCTGGTCTCGTGGGCTCGGAGATGTG | Thermo Fisher | CUT&Tag library prep |
| v2_Ad2.9_GCTACGCT | CAAGCAGAAGACGGCATACGAGATAGCGTAGCGTCTCGTGGGCTCGGAGATGTG | Thermo Fisher | CUT&Tag library prep |
| v2_Ad2.10_CGAGGCTG | CAAGCAGAAGACGGCATACGAGATCAGCCTCGGTCTCGTGGGCTCGGAGATGTG | Thermo Fisher | CUT&Tag library prep |
| v2_Ad2.11_AAGAGGCA | CAAGCAGAAGACGGCATACGAGATTGCCTCTTGTCTCGTGGGCTCGGAGATGTG | Thermo Fisher | CUT&Tag library prep |
| v2_Ad2.12_GTAGAGGA | CAAGCAGAAGACGGCATACGAGATTCCTCTACGTCTCGTGGGCTCGGAGATGTG | Thermo Fisher | CUT&Tag library prep |
| v2_Ad2.13_TGGATCTG | CAAGCAGAAGACGGCATACGAGATCAGATCCAGTCTCGTGGGCTCGGAGATGTG | Thermo Fisher | CUT&Tag library prep |
| v2_Ad2.14_CCGTTTGT | CAAGCAGAAGACGGCATACGAGATACAAACGGGTCTCGTGGGCTCGGAGATGTG | Thermo Fisher | CUT&Tag library prep |
| v2_Ad2.15_TGCTGGGT | CAAGCAGAAGACGGCATACGAGATACCCAGCAGTCTCGTGGGCTCGGAGATGTG | Thermo Fisher | CUT&Tag library prep |
| v2_Ad2.16_AGGTTGGG | CAAGCAGAAGACGGCATACGAGATCCCAACCTGTCTCGTGGGCTCGGAGATGTG | Thermo Fisher | CUT&Tag library prep |
| **HDR templates** | **Sequences** | **Company** | **Experiment** |
| crRNA - HA-tag | rCrU rUrCrU rCrArU rArGrA rUrArU rGrGrA rGrUrC rGrUrU rUrUrA rGrArG rCrUrA rUrGrC rU | IDT | HDR |
| HDR donor - HA-tag | C*T* GAT ACT GAG AAT TAA CAT TTT CCC TTC TCA TAG ATA TGT ACC CAT ATG ATG TTC CAG ATT ACG CTG AGT CTG GAG AAC GGT TAC CAT CCT CAG CAG CCT CCT CTA* C*T | IDT | HDR |
